# Systematic mapping of protease-substrate specificity through geometric graph learning

**DOI:** 10.64898/2026.04.08.717168

**Authors:** Xudong Guo, Zixu Ran, Yue Bi, Tong Pan, Heyun Sun, Yi Hao, Runchang Jia, Cong Wang, Qi Zhang, Lukasz Kurgan, Jiangning Song, Fuyi Li

## Abstract

Protease-substrate specificity is central to cellular regulation and disease pathogenesis, and accurately modelling its structural determinants remains challenging. Substrate recognition is governed by spatial constraints and higher-order relationships that extend beyond local sequence motifs. Most computational approaches rely predominantly on motif-centric or sequence-based representations, limiting their ability to capture the geometric and relational structure underlying enzymatic specificity. Here, we introduce OmniCleave, a structure-aware geometric graph learning framework for modelling protease-substrate specificity at scale. OmniCleave is trained on 57,278 structure-informed protease-substrate pairs derived from 9,651 substrates spanning over 100 proteases across six distinct families. The framework integrates multi-scale structural graphs with higher-order protease relational topology, explicitly encoding spatial context and inter-protease dependencies within a unified geometric representation. This formulation moves beyond local pattern recognition and enables transferable modelling across six protease families. Across large-scale benchmarks, the framework consistently outperforms existing approaches and reveals interpretable geometric determinants underlying substrate recognition. Experimental validation confirms three previously unannotated caspase-3 substrates and 21 cleavage sites predicted by OmniCleave, supporting the biological relevance of the learned representations. Together, OmniCleave provides a scalable geometric framework for modelling protease-substrate specificity, with practical utility for systematic analysis of protease biology.

## Introduction

Proteases are enzymes that hydrolyse peptide bonds in target proteins or peptide substrates, playing an essential role in processes such as cell cycle regulation, signalling, and protein degradation (1–4). Their functions are mediated through protease-substrate interactions, where proteases specifically recognise and cleave their substrates at defined positions. Cleavage typically occurs at the P1-P1’ site, where P1 denotes the residue on the N-terminal side of the scissile bond and P1’ the residue on the C-terminal side (5). Some proteases display substrate specificity, preferentially recognising particular residues or motifs within their targets, whereas others exhibit broader or less defined specificity (6). Accurately identifying protease substrate cleavage sites is therefore critical to understanding the diverse biological functions of proteolysis in both health and disease. Such insights have highlighted how proteolysis underpins fundamental aspects of physiology, shapes diverse cellular and tissue processes, and contributes broadly to human health and disease across many biological systems (2, 3, 7, 8).

Protease-specific substrate cleavage sites are typically identified by experimental assays such as proteomic peptide library (PPL) profiling and N-terminal COFRADIC (9, 10). These approaches are usually costly and do not scale across the vast protease repertoire. Large-scale protein databases contain a substantial number of protease-associated sequence records. UniProtKB (Release 2026_02) contains 2,663,559 protein entries annotated with the Protease keyword, of which only 12,286 are reviewed (11). Although protease-associated sequence records are abundant, experimentally supported annotations of protease-substrate relationships and cleavage-site specificity remain comparatively scarce, limiting the systematic investigation of protease-mediated processes.

Computational methods have been developed to address these limitations, which can be categorised into two groups: scoring function-based and machine learning-based (12). Based on relatively simple models, the scoring function-based methods use protein sequences to estimate the likelihood of cleavage at specific sites. Their use of simple sequence representations (features) constrains their predictive power. For instance, amino acid frequency (AAF), nearest neighbour similarity (NNS), position-specific scoring matrices (PSSM), and position weight matrices (PWM) are utilized by the scoring function-based tools such as PoPs (13), SitePrediction (14), PeptideCutter (15), and GPS-CCD (16). In contrast, machine learning-based methods rely on more sophisticated features, computed from protein sequences and their corresponding structural and functional characteristics, to learn predictive models from training data. A few representative machine learning-based tools include EM-HIV (17), TSignal (18), Cascleave (19, 20), PROSPER (21), CleavPredict (22), iProt-Sub (23), PROSPERous (24), Pripper (25), DeepCleave (26), Procleave (27), ProsperousPlus (28), and PGCN (29). These methods were developed by multiple independent research groups and collectively represent the major methodological advances in protease cleavage-site prediction, ranging from conventional feature-based classifiers to deep-learning and structure-informed models. These machine learning-based methods have been widely adopted by academia and industry (30, 31). Although several existing tools, such as Procleave and PGCN, have incorporated structural information, they generally focus on individual proteases without explicitly considering interactions among different proteases. As a result, these methods do not fully exploit protease-protease relationships to share information across proteases. Because proteases may be functionally connected through common pathways, molecular complexes, and protease-protease interactions, modelling them as isolated prediction tasks may overlook informative relational signals (32). A key unmet need is therefore a unified multi-protease framework that jointly learns cleavage specificity while integrating substrate sequence, structural context, and protease-protease relationships.

Recent advances in geometric deep learning provide robust support for integrating protein sequence and structural information across a variety of biological prediction tasks (29, 33–36). To this end, we present OmniCleave, a novel method that facilitates a systematic and comprehensive prediction of protease-substrate interactions. OmniCleave produces highly accurate predictions by applying an innovative structure-aware geometric graph transformer model and multi-view structural features of substrates that we generated using modern protein sequence embeddings (33, 37, 38). Moreover, our model incorporates a protease-protease interaction network that covers over 100 proteases, leveraging protease-protease interaction links as prior knowledge to predict substrate cleavage sites across the entire protease panel in a single inference step. Empirical tests demonstrate that OmniCleave is significantly more accurate than existing methods, especially in deciphering the many-to-one protease-substrate relationships. Moreover, our analyses reveal that OmniCleave learns substrate-specific structural preferences and captures the influence of inter-protease interactions on substrate cleavage site prediction and evolutionary signals across protease families.

## Results

### Overview of the OmniCleave framework

Unlike the current models that were designed for individual protease types, OmniCleave simultaneously predicts cleavage sites across diverse proteases by incorporating fine-grained inter-protease network information as relational priors, while jointly leveraging proteases and substrate sequences and structures. Fig. 1 shows the overall OmniCleave framework, which consists of four elements: data collection and pre-processing (Fig. 1a) and the hierarchical graph representation for substrate cleavage sites (Fig. 1b) that we collectively use to collect inputs to the OmniCleave model; the OmniCleave model that incorporates the protease-protease interaction network and a graph transformer-based protease-substrate predictor (Fig. 1c); and example applications for the outputs generated by OmniCleave (Fig. 1d).

**Figure 1.**
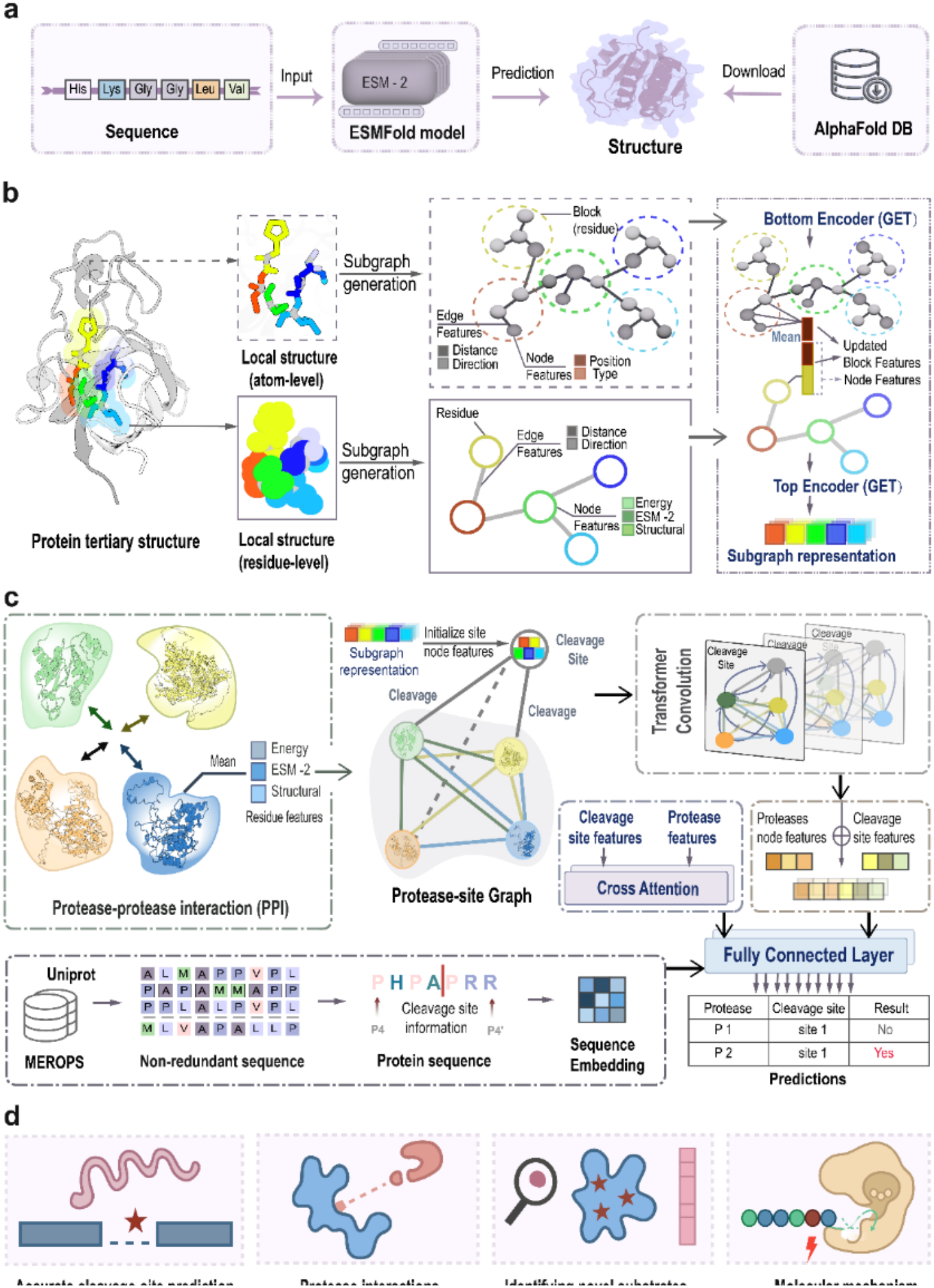
Overview of the OmniCleave framework. **a.** Collection of 3D structures of substrates. **b**. OmniCleave represents 3D structural information of substrates using hierarchical graph representations at atomic and residue levels centred on cleavage sites. **c**. OmniCleave integrates protease-protease relationships via a PPI-based graph neural network and models protease-substrate interactions with a graph transformer. This unified framework integrates proteases and substrates into a shared graph space, enabling flexible and large-scale prediction of substrate cleavage sites across diverse proteases. **d**. Applications of OmniCleave’s outputs include substrate cleavage site prediction, characterization of protease-protease interactions, discovery of novel protease substrates, and mechanistic studies of protease-mediated catalysis.

We curated the datasets used for model training and evaluation, which contain 57,278 substrate cleavage sites (Supplementary Table S1), from MEROPS(39) and UniProt(40) databases, along with their corresponding 3D structures that we retrieved from the AlphaFold database(41, 42) and generated with ESMFold(43) (Fig. 1a). These cleavage sites are associated with 103 proteases across six protease superfamilies, including Aspartic, Cysteine, Glutamic, Metallopeptidase, Asparagine, and Serine(39). To construct a balanced benchmark, we also included 57,278 randomly sampled non-cleavage sites. As shown in Fig. 1b, the datasets described above, together with information on protease-protease relationships, are represented as a heterogeneous graph at hierarchical scales (atomic and amino acid levels), which serves as input to the OmniCleave model.

OmniCleave is a deep learning architecture that consists of three major components: (i) A cleavage-centric hierarchical encoder, which is implemented as a hierarchical graph network that enables information exchange across the residue and atom level scales for the substrate protein. As illustrated in Fig. 1b, a hierarchical subgraph of the substrate protein is constructed around the cleavage site, guided by the putative 3D structure of the substrate at both residue and atomic resolutions. The residue-level subgraph covers residues with Cα-Cα distances ≤10 Å from the cleavage site, and each residue is annotated with Rosetta energy terms, ESM-2 embeddings, and secondary structure information. At the atomic level, subgraphs enrich the atomic-level representations with atom types and their spatial arrangement. For both levels, edge features combine distance encodings with directional vectors. Together, these residue- and atomic-level details produce complementary representations of the cleavage-site microenvironments, providing information-rich input for the OmiCleave predictive model.

Targeting the vicinity of the cleavage sites focuses on functionally critical regions while reducing computational overhead that would be induced by enlarging this area. (ii) A protease-protease interaction (PPI) network module that constructs a PPI network, as shown in Fig. 1c. The relational priors derived from the PPI network are incorporated through a graph neural network in which each protease is represented as a node, and edges denote protease-protease interactions annotated using the STRING database (44). This design facilitates capturing cooperative patterns between proteases, providing an important cellular-level context that strengthens predictions of the substrate cleavage sites when compared to modelling each protease individually. (iii) The graph transformer-based protease-substrate interaction module (Fig. 1c), which embeds proteases and substrate cleavage-site subgraphs into a unified heterogeneous graph utilizing multiple transformer convolutional layers, where each cleavage site is connected to its interacting proteases. By propagating messages across protease-protease and protease-cleavage site links, this module combines broader/protease context and local/cleavage site signatures, thereby producing fused protease-aware representations. Through an iterative training process that involves a message-passing mechanism, these representations are subsequently projected by attention-based layers into scalar probability values that denote the likelihood of a cleavage site.

### OmniCleave accurately predicts protease substrate cleavage sites

We comprehensively benchmarked OmniCleave on the large dataset with 103 protease substrate cleavage site datasets that span six protease superfamilies (Supplementary Table S2). We compared OmniCleave with six state-of-the-art (SOTA) computational methods, including Procleave (27), PROSPERous (24), DeepCleave (26), SitePrediction (14), PeptideCutter (15), and ProsperousPlus (28). All methods were evaluated on the same independent test set. However, as the training data for some comparison tools are unavailable, it is unclear whether their training sets included the substrate proteins present in our test set. This potential data overlap may have given those tools an advantage in the comparison. We assessed predictive performance using the popular area under the receiver operating curve (AUC), the area under the precision-recall curve (AUPR), and the F1 score. OmniCleave achieved AUCs above 0.9 for 48 proteases and above 0.8 for 75 proteases (Fig. 2a). Furthermore, under a more stringent protein similarity threshold (< 30%), OmniCleave maintained AUC values greater than 0.9 for 58 proteases and greater than 0.8 for 74 proteases (Fig. 2f). For a collection of seven arguably key proteases, including caspase-3 (C14.003), caspase-7 (C14.004), MMP-2 (M10.003), MMP-9 (M10.004), MMP-7 (M10.008), granzyme B (S01.010), and thrombin (S01.217), OmniCleave consistently outperformed the six SOTA methods (Fig. 2b). Both OmniCleave and Procleave outperformed the five sequence-based methods (ProsperousPlus, PROSPERous, DeepCleave, SitePrediction, and PeptideCutter), underscoring the benefit of integrating putative structure information.

**Figure 2.**
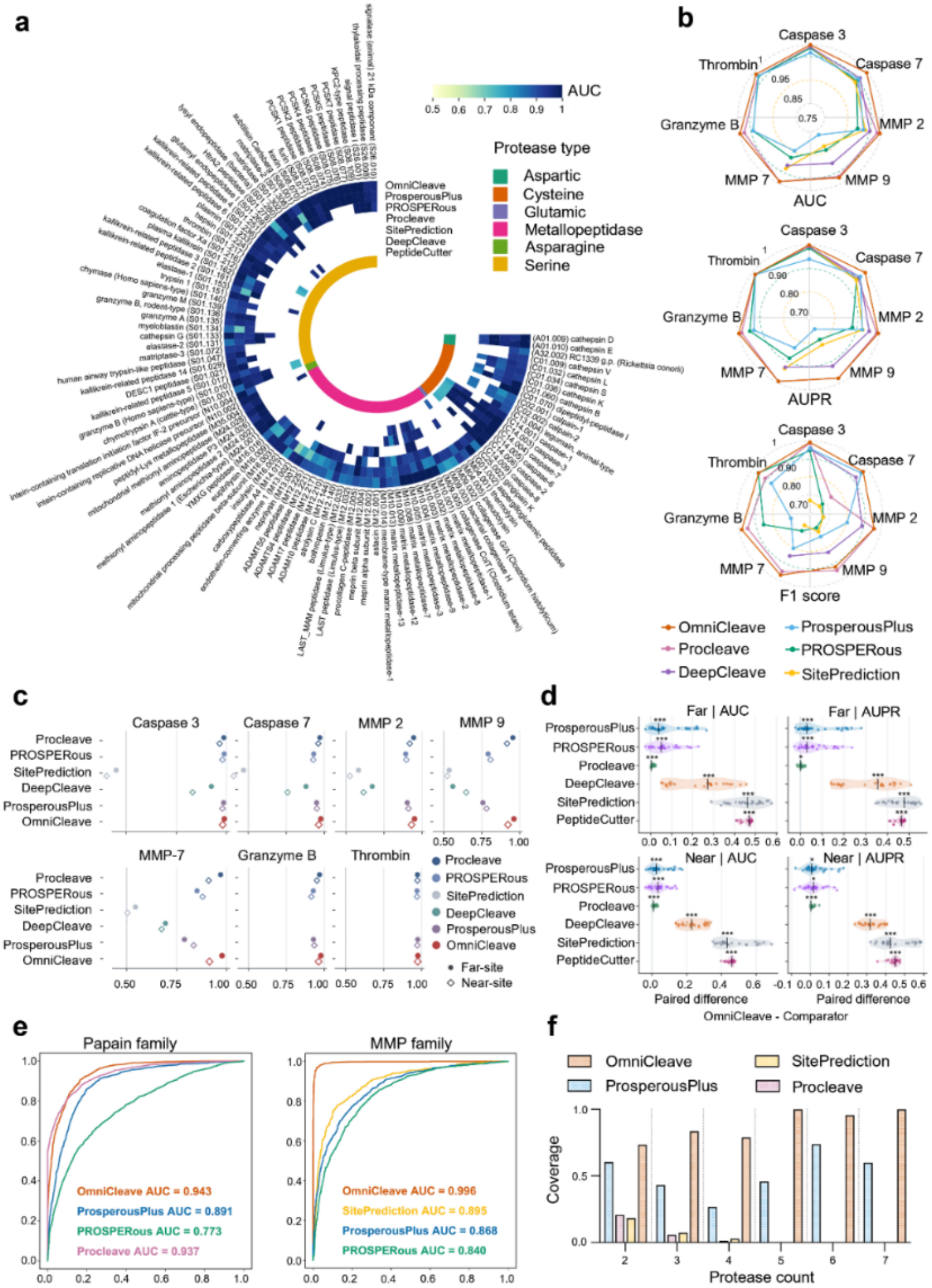
Benchmarking performance of OmniCleave. **a.** Comparison of AUC values across 103 protease substrate cleavage datasets between OmniCleave and six representative state-of-the-art tools: ProsperousPlus, PROSPERous, Procleave, SitePrediction, DeepCleave, and PeptideCutter. White cells indicate proteases that are not supported by a given method. **b**. Performance comparison of OmniCleave and the five methods on seven representative protease datasets based on the AUC, AUPR, and F1-score metrics. **c**. Performance comparison of OmniCleave and the five methods on seven representative protease datasets (far- and near-site) based on the AUC. **d**. Pairwise differences between OmniCleave and other methods on far- and near-site datasets of seven proteases. The vertical black lines indicate the medians of the differences for the respective tools, and the asterisks denote statistical significance. ***: q < 0.001; **: q < 0.01; *: q < 0.05. **e**. ROC curves of OmniCleave and the four methods across the Papain and Matrix Metalloproteinases (MMP) protease families. **f**. Performance comparison of OmniCleave and the three methods in many-to-one protease-substrate site identification.

To evaluate the effect of negative sampling on model performance, we constructed two additional datasets using far-site and near-site negatives (see Methods). OmniCleave was then compared wih six existing tools on both datasets (Fig. 2c). The results demonstrated that OmniCleave maintained a robust performance advantage under both settings. We further conducted five independent rounds of negative sampling, model training and testing for each strategy. Paired statistical comparisons were performed across seven representative proteases (Fig. 2d). For each protease and experimental run, we calculated the performance difference between OmniCleave and each competing tool. For both AUC and AUPR, most paired differences were positive, indicating that OmniCleave generally outperformed the other methods under both negative-sample definitions. Compared with DeepCleave, SitePrediction, and PeptideCutter, OmniCleave showed larger and more consistently distributed performance gains on both the far-site and near-site datasets. The differences relative to ProsperousPlus, Procleave, and PROSPERous were smaller, but still generally favoured OmniCleave. We used one-sided Wilcoxon signed-rank tests and applied the Benjamini–Hochberg correction for multiple comparisons. Most pairwise comparisons remained statistically significant after correction. Taken together, these results indicate that OmniCleave’s performance advantage is not limited to a particular negative-sampling strategy and remains consistent across independent experimental runs.

We compared average AUCs across four protease families (Papain, MMP, Caspase, and Chymotrypsin), which are covered by four SOTA tools (Fig. 2e and 2g). OmniCleave consistently outperforms these four tools across the four protease families, with particularly strong performance for MMP and Chymotrypsin. OmniCleave also substantially improves over the structure-based Procleave (Fig. 2b). OmniCleave consistently achieved leading predictive performance across the Cysteine, Metallopeptidase, and Serine protease superfamilies (protease type), demonstrating strong generalizability across diverse protease superfamilies (Fig. 2h). It is worth noting that some existing tools support only a limited number of proteases. The remaining three superfamilies were not included in this cross-tool comparison because they contained too few supported proteases to yield representative and statistically stable family-level aggregate results. The t-SNE visualization of the model-learned embeddings across the seven representative proteases (extended Fig. 2i) reveals that the cleavage samples (positives) and non-cleavage samples (negatives) show clear separation, in most cases forming distinct and well-defined clusters. The underlying ability to effectively capture distinct and protease-substrate-specific signatures explains why OmniCleave produces very accurate results.

Protease-substrate interactions are not isolated events but instead occur in a context of a complex regulatory network. Within this network, some proteases form one-to-one relationships with their substrates. However, our analysis reveals a prevalence of many-to-one protease-substrate interactions. For instance, using data from the MEROPS database, a substantial number of substrates were recognised and cleaved by multiple distinct proteases at the same sites. In extreme cases, a single cleavage site was targeted by up to 20 different proteases (see Supplementary Tables S3-S4). We assessed the capability of OmniCleave to uncover these relationships by constructing and using a many-to-one protease-substrate cleavage dataset sampled from the test set. DeepCleave was excluded from this comparative experiment due to its insufficient protease coverage. As shown in Fig. 2f, OmniCleave consistently achieved markedly higher coverage across all protease-count categories compared with the three SOTA methods (ProsperousPlus, PeptideCutter, and Procleave). Its advantage was particularly evident when substrates were targeted by a higher number of proteases (≥5), where OmniCleave maintained near-complete coverage, while the performance of the other methods dropped substantially. Notably, in cases involving multiple proteases acting on the same substrate, OmniCleave demonstrated exceptional sensitivity and generalisation capacity. This advantage over the other tools can be explained by the fact that OmniCleave utilises the PPI networks that represent the many-to-one interactions. Taken together, these findings indicate that OmniCleave produces accurate results for the cleavage sites in one-to-one and many-to-one interaction scenarios. An exploratory comparison with AlphaFold 3, together with its methodological limitations, is provided in Fig. 2j and the Supplementary Note.

Proteolytic cleavage sites may occur in flexible regions or regions with uncertain structures. We therefore examined whether the structural confidence of predicted protein models affected OmniCleave’s performance. First, we compared the P1-residue pLDDT distributions between cleavage and putative negative sites (non-cleavage sites) with most samples located in high-confidence regions (Fig. 2k). Different negative-sampling strategies slightly changed the pLDDT distributions of putative negative sites. However, no clear difference in pLDDT was observed that could readily separate positive and negative samples. Fig. 2l further demonstrates that OmniCleave maintains stable predictive ability across different pLDDT intervals. In the random-negative dataset, the average AUC for all three confidence groups was approximately 0.92, and the average MCC was approximately 0.68-0.70. For far-site dataset, AUC values ranged from 0.885 to 0.915, with MCC values between 0.602 and 0.663. Similar results were observed for the near-site dataset, where AUC ranged from 0.902 to 0.924 and MCC ranged from 0.650 to 0.709. Notably, OmniCleave achieved an AUC of at least 0.90 even for sites with pLDDT below 70. These findings suggest that the model maintains good predictive performance across different levels of structural confidence and does not rely solely on high-confidence structural regions. We also examined the relative solvent accessibility (ASA) of P1 residues (Fig. 2m). Cleavage sites was generally slightly higher than that of non-cleavage sites. This is consistent with the biophysical expectation that protease cleavage sites should, on average, be more exposed to solvent compared to a selection of randomised residues. However, the ASA distributions of the two groups showed considerable overlap and could not clearly distinguish cleavage from non-cleavage sites. The negative samples in the far-site and near-site datasets also showed similar ASA distributions, suggesting that these sampling strategies did not introduce substantial differences in solvent accessibility. Therefore, ASA may be an effective structural feature for cleavage, but it cannot explain the model’s classification performance on its own. Figure 2n further shows that the secondary structure composition of cleavage sites and non-cleavage sites is generally similar. The proportions of coil, helix, and strand were generally comparable between cleavage and putative negative sites. Only small differences were observed in the far-site and near-site datasets.

Overall, the choice of negative-sampling strategy had a modest effect on model performance. OmniCleave achieved AUC values of 0.924, 0.898, and 0.915 for the random-, far-site-, and near-site-negative datasets, respectively. The corresponding MCC values were 0.695, 0.629, and 0.678. OmniCleave still maintains high predictive performance under the more stringent far-site and near-site negative sampling settings. Combining pLDDT stratification results with ASA and secondary-structure distribution analyses supports the model’s robustness to the definition of the putative negative samples and reduces the likelihood that the model will classify based solely on simple biases, such as low-confidence structures or specific secondary structures.

### OmniCleave captures the structural contexts of cleavage events

The ability to capture the structural context of cleavage events at both atomic and residue levels is essential to facilitate accurate modelling/prediction of these interactions. To demonstrate that, we compared OmniCleave, which utilises the hierarchical subgraphs that model cleavage sites at both levels, with a variant of our model that is restricted to the residue-level information (Fig. 3a). OmniCleave consistently outperforms the residue-level variant for the cleavage-site prediction across the seven representative proteases and across the AUC, AUPR, and F1-score metrics. These results underscore the need to integrate both residual and atomic-level contexts. Moreover, we observed distinct structural distributions between cleavage sites (positives) and non-cleavage sites (negatives) (Fig. 3b), with the non-cleavage sites showing more dispersed subgraph patterns.

**Figure 3.**
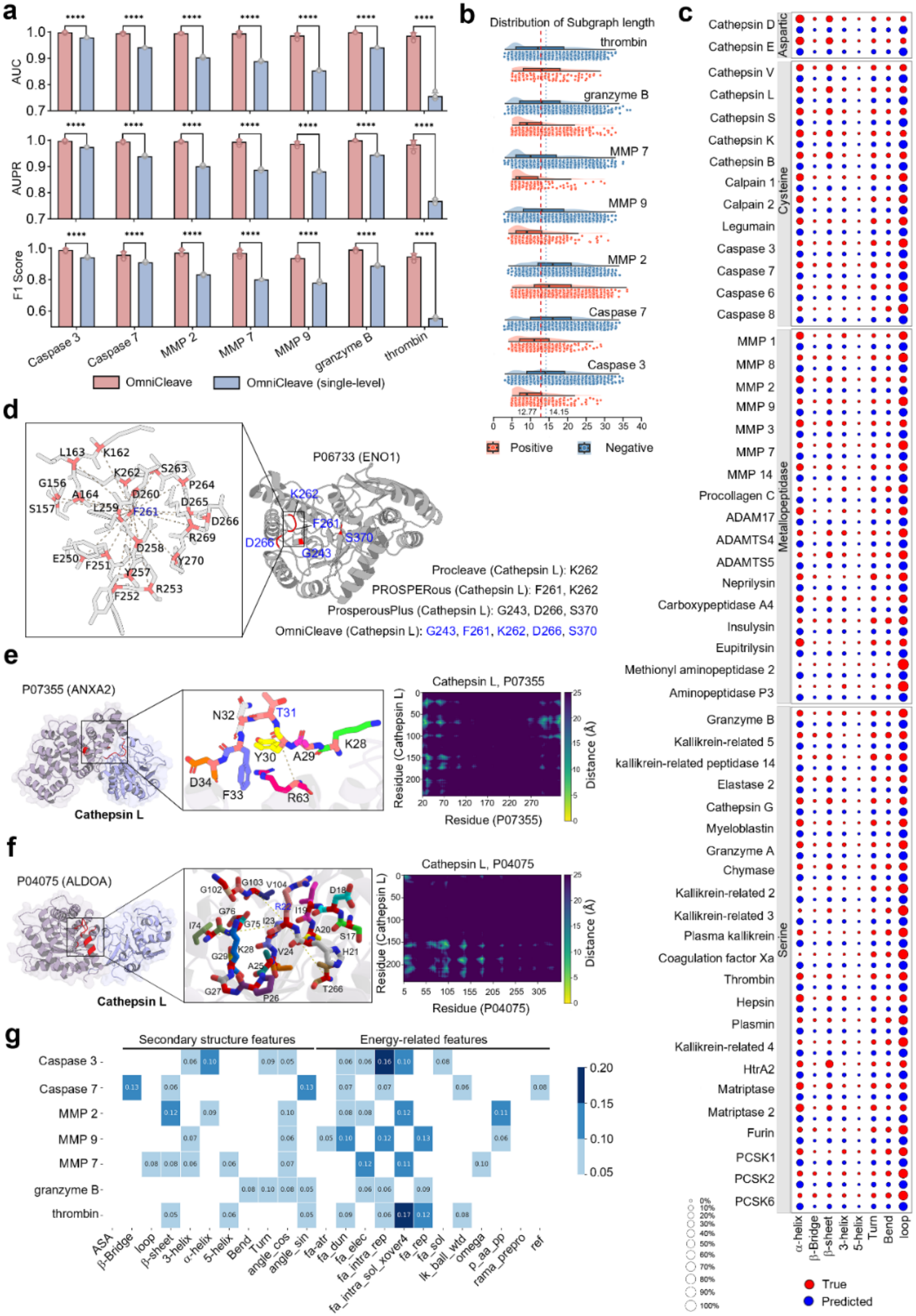
Mechanistic insights into protease-substrate cleavage via hierarchical graph representation. **a.** Performance comparison between OmniCleave and the residue level-only variant. b. Distribution of subgraph sizes for cleavage-centred (positives) versus non-cleavage-centred (negatives) subgraphs. **c**. Predicted versus observed secondary-structure states of P1 residues across 54 human proteases; true distributions in red, OmniCleave predictions in blue; dot size indicates frequency. **d**. Improvement in cleavage-site recognition enabled by local subgraph representation for Cathepsin L-P06733 complex. **e**. Structural visualization of the Cathepsin L-ANXA2 (P07355) complex and residue distributions within cleavage-centred subgraphs. **f**. Structural visualization of the Cathepsin L-ALDOA (P04075) complex and residue distributions within cleavage-centred subgraphs. **g**. Feature perturbation heatmap highlighting the dominant contributions of secondary structure and Rosetta energy features to OmniCleave predictions.

We analysed whether OmniCleave captures the correct secondary-structure context for its predicted cleavage sites. More specifically, we compared the secondary-structure states of P1 residues at the predicted cleavage sites against their native structures across 54 human proteases (Fig. 3c). We categorised residues into eight secondary structure states: α-helix, β-bridge, β-sheet, 3-helix, 5-helix, turn, bend, and loop. OmniCleave’s predictions closely mirrored the true distributions of these secondary structure states, with cleavage sites most frequently located in loops, α-helices, β-sheets, turns, and bends. Loops are the most common, which is consistent with their flexibility and accessibility to enzymatic recognition, whereas β-bridges are rare and 5-helices are virtually absent(19, 45). Distinct secondary structure distribution patterns were also observed across protease families: Aspartic proteases (e.g., Cathepsin D/E) are enriched in α-helices, loops, and β-sheets. Cysteine proteases displayed two distinct trends: Cathepsins V/L/S/K/B are enriched in α-helices, β-sheets, and loops, whereas Caspases 3/6/7/8 are predominantly associated with loops, α-helices, turns, and bends. Metallopeptidases (e.g., MMP1/2/3/7/8/9) exhibit broad distributions across α-helices and loops, with secondary enrichment in β-sheets and turns. Serine proteases (e.g., Plasmin, Furin, Granzyme A, Thrombin) are largely concentrated in loops and α-helices. For certain proteases, such as Coagulation Factor Xa, the predicted distribution deviates from the true distribution, likely due to the limited sample size, which consists of only 24 instances.

Next, we provide more detailed insights into the advantages of OmiCleave’s predictions that rely on enriched structural representations of cleavage sites, when compared to the results generated by SOTA sequence-based algorithms. We use cathepsin L (C01.032, PDB 2xu3) and its substrate P06733 as an illustrative example. For sequence-based algorithms, the input consists solely of the substrate protein’s amino acid sequence, from which these methods automatically extract local sequence windows of varying lengths upstream and downstream of the putative cleavage site. In contrast, OmniCleave operates on sequence information combined with the substrate protein’s structure, capturing three-dimensional structural features from all residues located within a 10Å spatial radius of the candidate site. OmniCleave accurately identified all five of its experimentally verified cleavage sites (G243, F261, K262, D266, and S370), whereas sequence-based ProsperousPlus, PROSPERous, and Procleave predicted three, two, and only one site, respectively (Fig. 3d). Notably, at the K262 site, Procleave relied on the information concerning only the eight flanking residues, while OmniCleave integrated a broader local substructure encompassing 20 residues, including both P4-P4’ and distal positions. The use of the richer structural representation explains the corresponding substantial difference in performance. In the case of cathepsin L with the P07355 substrate, OmniCleave accurately identified the T31 cleavage site and captured its interaction with the sequence-distant residue R63 (Fig. 3e). For P04075, it also detected R22 cleavage by integrating context from both proximal residues (e.g., H21, I23, and V24) and distal residues (e.g., G754, G103, and T266) (Fig. 3f). Contact maps confirmed that these distal residues lie in the protease-substrate interface, underscoring their contribution to cleavage recognition.

To evaluate the contribution of node features within residue-level subgraphs, we performed random perturbations and measured the resulting changes in predictions(29) (Materials and Methods; Supplementary Tables S5-S6). Across seven representative protease datasets, energy-related features, including van der Waals interactions and backbone constraints, were the most influential (Fig. 3g). Features, including β-bridges, loops, helices, bends, turns, and angles, consistently exerted non-negligible effects across protease datasets, emphasising the importance of local secondary structure in substrate recognition.

### Protease-protease interaction data used by OmiCleave drives accurate multi-protease cleavage prediction

OmniCleave incorporates PPI networks to improve substrate recognition across diverse collections of proteases, including the many-to-one protease-substrate interactions. Using STRING (44), we constructed protease-protease interaction (PPI) networks for 27 species, with the human network being the largest and most connected, encompassing 54 proteases. We further applied Metascape (46) and Cytoscape (47) to perform MCODE clustering and visualization (Fig. 4a). MCODE clustering (48) of the human network identified four functional modules (MCODE1-4) with distinct cooperative patterns (Fig. 4e). MCODE1 (CTSS, CTSV, CTSK, CTSL, MMP3, MMP7, MMP9), enriched for cathepsins and matrix metalloproteinases, was linked to collagen catabolism and metallopeptidase activity, suggesting roles in extracellular matrix (ECM) remodelling. MCODE2 (CASP3, CASP6, CASP7, CASP8, CAPN2), comprising caspases and calpain, was associated with protein processing and cysteine-type endopeptidase activity, consistent with apoptotic signalling. MCODE3 (CMA1, CTSG, ELANE, KLKB1, PLG), including neutrophil serine and fibrinolytic proteases, was enriched for ECM disassembly and serine hydrolase activity, implicating roles in inflammation and tissue repair. MCODE4 (F10, F2, HTRA2), containing coagulation factors and HTRA2, was localised to membranes and linked to serine-type endopeptidase activity, highlighting functions in coagulation and extracellular signalling. Together, these results justify our use of the protease interaction information encoded in the PPI network in the OmniCleave model. By constructing additional nodes and integrating substrate nodes into PPI, information on proteases and substrates is exchanged between nodes in our model via a message-passing mechanism. Graph embeddings fusing protease and substrate information, as well as embeddings of substrate and protease nodes, are then used to predict protease-substrate relationships. This information exchange enables OmniCleave to capture cooperative cleavage patterns among functionally related proteases and to analyse many-to-one protease-substrate relationships.

**Figure 4.**
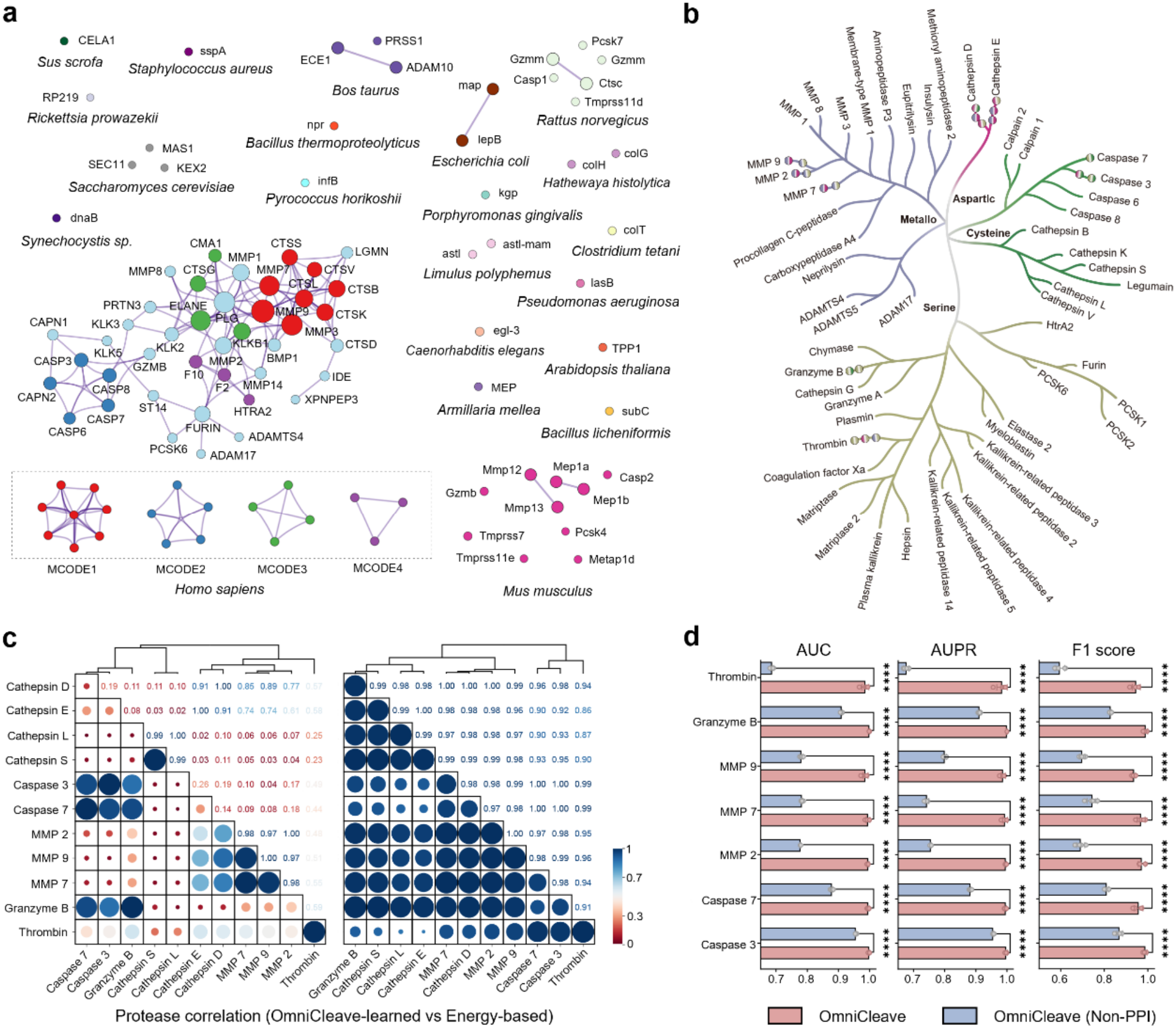
Protease-protease interactions and feature correlations shape predictive model performance. **a.** Protease-protease interaction networks across species, constructed from STRING. The human protease subnetwork is the largest and most densely connected. Four functional modules (MCODE1-4) were identified through MCODE analysis and are highlighted below. **b**. Phylogenetic tree of 54 human proteases based on catalytic domain sequences. Catalytic types are colour-coded; bi-colored spheres indicate proteases with high correlation (Pearson correlation), with colours at each end representing their respective types. **c**. Correlation heatmap of 11 proteases using model-updated features (left) or original energy-term features(right). **d**. Performance comparison of OmniCleave with or without the PPI module on seven test-set proteases, evaluated by AUC, AUPR, and F1 score.

The PPI network also reflects evolutionary relationships among proteases. We constructed a phylogenetic tree of 54 human proteases based on their catalytic domain sequences (Fig. 4b). The resulting tree topology closely recapitulates the MEROPS classification (39) based on catalytic type, with proteases segregating into clearly defined clans (AA, CA, CD, MA, MG, SB, PA; Supplementary Table S2) corresponding to a common evolutionary origin. Aspartic proteases (Cathepsin D/E) form a single branch, while Cysteine proteases split into two major branches: Cathepsin B/K/S/L/V with Legumain, and Calpain1/2 with Caspase3/6/7/8. Among metalloproteases, Neprilysin clusters with ADAMTS4/5 and ADAM17, whereas MMPs (MMP1/2/3/7/8/9) form a separate branch. Serine proteases have multiple branches, including a distinct cluster of kallikrein-related peptidases (KLK2/3/4/5/14). Integration of protease correlation patterns (Fig. 4c) further uncovers strong associations across evolutionary branches, such as between MMP2/7/9 and Aspartic proteases or Thrombin, despite their considerable evolutionary distance (Fig. 4b). These findings suggest that, whereas the phylogenetic tree primarily reflects conservation of catalytic domains, integrating PPI networks with substrate specificity enables OmniCleave to reveal novel functional associations that span evolutionary lineages.

To investigate whether OmniCleave captures inter-protease relationships, we computed Pearson correlations between its learned protease feature vectors and, for comparison, between raw Rosetta(49) energy terms (Fig. 4c). Given the successful applications of Rosetta energy terms in protein structure prediction and protease design(29), we used them as the reference characterizations of proteases. Energy-based correlation analysis showed that proteases within the same family have highly similar energy characteristics, while proteases from different families exhibit certain differences. In terms of OmniCleave-learned correlation analysis, strong correlations were observed within protease families, such as Caspase 3-7 and MMP7-9, while notable cross-family correlations were also detected. For instance, between Granzyme and Caspase 7, or MMP9 and Cathepsin D, indicating that OmniCleave can propagate substrate information across proteases (Fig. 4b). To further quantify the contribution of protease-protease interaction information, we compared the full OmniCleave model with a variant that removed the PPI interaction module, while keeping all substrate features, training settings, and datasets unchanged. As shown in Figure 4d, removing PPI module consistently reduced the prediction performance of OmniCleave. Clear decreases in AUC, AUPR, and F1 scores were observed across all seven representative proteases. This performance degradation was not limited to a single protease family but was observed across all seven proteases, indicating that the interaction network provides complementary information for multi-protease cleavage prediction. Importantly, although the model without PPI still achieved relatively high performance, it consistently performed worse than the complete model. These results suggest that substrate sequence and structural features provide a basis for cleavage prediction, while PPI information further improves the model by introducing additional protease-specific context.

### Sequence and structural representations provide complementary information

We expanded the ablation analysis to assess how strongly OmniCleave relies on local P4-P4’ sequence motifs. We also evaluated the contributions of individual sequence and structural components (Supplementary Material). Five variants were considered: w/o GNN (GET), w/o DSSP, w/o energy, w/o seq-based, and Seq + PPI. In the w/o GNN (GET) variant, the GET encoders applied to the local residue-level and atom-level substrate graphs were removed, whereas the P4-P4’ sequence-derived representations, DSSP descriptors (DSSP-based features), Rosetta-derived residue-level score components (energy-based features), and PPI information were retained. The w/o DSSP and w/o energy variants selectively removed the DSSP-based features and energy-based features, respectively, while retaining the remaining model inputs. In contrast, the w/o seq-based variant removed the P4-P4’ sequence-derived representations (chemical group feature CR and the amino acid (AA) type feature SR), while retaining the structural, energetic, and protease-relation features.

We further constructed a Seq + PPI variant that retained only the P4-P4’ sequence-derived substrate features S^R^ and C^R,^ together with the PPI-based protease representation. All substrate graph-based structural, DSSP-derived, and Rosetta-derived energy features were removed. The PPI representation was retained because OmniCleave jointly models 103 proteases within a single framework. Protease-specific conditioning is therefore needed to distinguish their cleavage preferences. Without protease-specific information, identical or highly similar P4-P4’ substrate windows could have different protease-specific cleavage labels. This is because different proteases may share cleavage motifs or target the same substrate positions. Thus, the Seq + PPI variant was designed to assess the predictive capacity of local substrate sequence information while preserving the protease context required for unified multi-protease prediction.

Across seven representative proteases and five independent runs (Table 1), the complete OmniCleave model achieved macro-averaged AUC, AUPR, and F1-score values of 0.9938, 0.9934, and 0.9659, respectively. The w/o GNN model achieved corresponding values of 0.9272, 0.9236, and 0.8726. Removal of the DSSP features resulted in values of 0.9102, 0.9044, and 0.8440, whereas removal of the Rosetta-derived energy descriptors resulted in values of 0.9167, 0.9112, and 0.8528. The Seq + PPI variant achieved macro-averaged AUC, AUPR, and F1-score values of 0.9217, 0.9240, and 0.8654, respectively. Although the sequence-focused model retained strong predictive ability, the complete OmniCleave model improved AUC, AUPR, and F1-score by 0.0721, 0.0694, and 0.1005, respectively. The complete model outperformed each structure-related ablation variant across all seven proteases, suggesting that graph-based, secondary-structure, and energetic representations provide complementary and impactful information. The largest reduction was observed after removing the P4-P4’ sequence-derived features. The w/o seq-based model achieved macro-averaged AUC, AUPR, and F1-score values of 0.6885, 0.6975, and 0.6218, respectively. These findings indicate that local sequence specificity is a major component of the predictive signal. Conversely, the Seq + PPI experiment shows that local sequence information, even when combined with PPI-based protease conditioning, is not sufficient to reproduce the performance of the complete model. The complementary results of the Seq + PPI and w/o seq-based variants indicate that sequence and structural information are not interchangeable. Local P4-P4’ sequence specificity provides a major predictive foundation, while graph-based structural, secondary-structure, energetic, and protease-context representations provide additional information.

**Table 1:** Comparison of ablation of different components in OmniCleave. The results represent the macro-averages derived from five independent experiments involving seven representative proteases. Seq + PPI retains only the P4-P4’ sequence-derived substrate representations (SR and CR) together with the PPI-based protease representation, while excluding residue- and atom-level graph encodings, DSSP-derived descriptors, and Rosetta-derived energy features. PPI information was retained to preserve protease-specific conditioning in the unified multi-protease framework.

| Model | AUC (Macro-average) | AUPR (Macro-average) | F1 score (Macro-average) |
|---|---|---|---|
| OmniCleave | 0.9938 | 0.9934 | 0.9659 |
| w/o GNN (GET) | 0.9272 | 0.9236 | 0.8726 |
| w/o DSSP | 0.9102 | 0.9044 | 0.8440 |
| w/o Energy | 0.9167 | 0.9112 | 0.8528 |
| w/o Seq-based | 0.6884 | 0.6975 | 0.6218 |
| Seq+PPI | 0.9217 | 0.9240 | 0.8654 |

We further quantified exact P4-P4’ motif overlap between the training and test sets (random split). The training and test sets contained 68,607 and 30,672 unique motifs, respectively, with only 1,004 exact motifs shared between the two partitions. These shared motifs accounted for 1.46% of the unique training motifs and 3.27% of the unique test motifs. Among positive samples, 553 motifs appeared as positive cleavage motifs in both partitions. These accounted for 1.71% of the 32,299 unique positive training motifs and 3.81% of the 14,508 unique positive test motifs. Thus, the large majority of positive motifs in the independent test set had not appeared as identically labelled positive motifs during training. Together with the substrate-level and 30% sequence-cluster-level separation, these results reduce the likelihood that the observed performance was driven primarily by direct memorisation of identical cleavage motifs.

Overall, the ablation results confirm that local P4-P4’ sequence preferences provide a major source of predictive information for OmniCleave. However, the low overlap of identical positive motifs between the training and test sets suggests that model performance is unlikely to result primarily from direct motif memorisation. Moreover, the higher performance of the complete model compared with the Seq + PPI variant supports the complementary contribution of substrate structural and contextual information beyond the local sequence window.

### OmniCleave identifies candidate protease substrates and associated biological processes

To explore the biological functions associated with protease substrates, we performed systematic GO and pathway enrichment analyses. These analyses were conducted on experimentally validated and OmniCleave-predicted substrates across 12 protease families. They covered GOBP, GOCC, GOMF, and KEGG (Fig. 5). The aim was to examine the functional profiles of these proteins rather than to validate cleavage at the predicted sites. By examining enriched terms associated with the actual substrates of proteases within each family, we observed functional convergence among several families, notably Caspase, MMP, Pepsin, and Legumain (Fig.5a). Among the 54 human proteases, significant functional convergence was observed among the substrates of Cathepsin D/E/V/L/S/K/B, Legumain, Caspase-1/3/6/7/8, and MMP-2/3 (Fig.5g). For Caspase family, the five considered caspase proteases (Caspase-1/3/7/6/8) exhibited varying degrees of substrate overlap (Fig. 5b). Caspase-1 and Caspase-3 shared 29 substrate proteins, while Caspase-3 and Caspase-7 shared 96. Across all five caspases, six substrates were common. Analysis of the top 10 enriched terms for these five proteases further revealed high functional convergence: 43 GO terms were associated with at least two proteases. Caspase-1 and Caspase-3 were linked to 26 terms each, whereas Caspase-7, Caspase-6, and Caspase-8 were associated with 15, 19, and 22 terms, respectively. Caspase-1 shared 11, 10, 15, and 7 enriched terms with Caspase-3, Caspase-7, Caspase-6, and Caspase-8. Five GO terms were shared among Caspase-1, Caspase-3, and Caspase-7, and one term appeared across all five proteases. For the MMP family, MMP-2 and MMP-3 share more substrates than other MMP proteases (Fig. 5g). Among the top 10 terms, the substrates of the seven MMP proteases share 26 enriched terms. Specifically, MMP-1/2/3/7/8/9/14 are enriched for 17, 5, 5, 14, 23, 19, and 16 terms, respectively; MMP-8 and MMP-9 share 18 terms; MMP-1/8/9 share 15 terms; MMP-1/8/9/14 share 8 terms; and MMP-1/8/9/7/14 share 4 terms (Fig. 5h). These results show that substrates of several proteases share similar functional annotations, particularly within the same protease family. The presence of numerous substrate overlaps within the same family reflects conserved substrate recognition rules within the family and suggests genetic redundancy (50, 51), as well as functional compensation within proteases of the same family (52, 53). This interconnected substrate landscape supports the integration of PPI networks into OmniCleave’s predictive framework.

**Figure 5.**
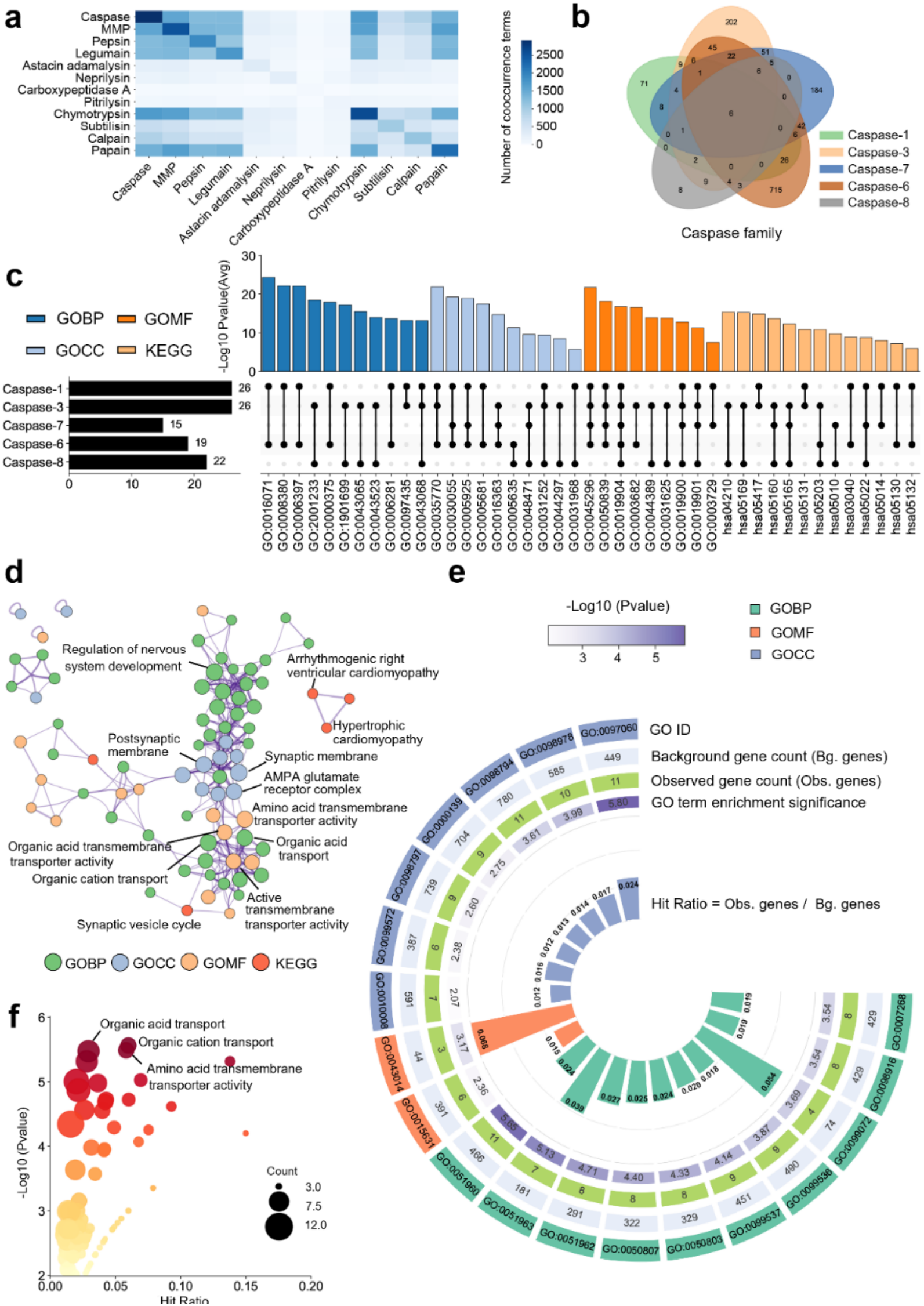
GO/KEGG enrichment for the experimentally validated and model-predicted substrates. **a.** Co-occurrence enrichment terms of 12 families of protease substrates. **b**. Distribution of shared substrates among 5 Caspase proteases. **c**. Distribution of shared enrichment terms among 5 Caspase proteases. **d**. Networks of enriched GO terms (BP, CC, MF) and KEGG pathways for predicted substrates of Caspase-3. **e**. Enrichment terms shared by experimentally validated and model-predicted substrates. **f**. Novel terms for model-predicted substrates.

We further investigated whether OmniCleave could be used to prioritize previously unannotated human protease substrates and explore their potential functional contexts. Tertiary structural models for human proteins were collected from the AlphaFold Protein Structure Database(42). To focus the analysis on previously unannotated candidates, we excluded all proteins containing at least one cleavage-site annotation in MEROPS Release 12.4 (as well as those in the training and test sets), irrespective of the annotated protease. Proteins without an available structural model were also excluded. After filtering, 14,332 human proteins were retained for proteome-scale prediction. We applied OmniCleave to these proteins for the 50 human proteases represented in our dataset. Protease-specific cleavage probabilities were calculated for all eligible residue positions, and residues with a predicted probability greater than or equal to 0.99 were retained as candidate cleavage sites. A protein was selected as a candidate substrate if at least one site had a predicted cleavage probability greater than 0.99 for any of the 50 proteases. Using this criterion, OmniCleave identified 1,379 unique human proteins as previously unannotated candidate substrates (Supplementary Table S7). These predictions should be interpreted as computationally prioritised candidates rather than experimentally validated substrates.

Using Caspase-3 as a representative example, we conducted an enrichment analysis on its 119 predicted substrates (Fig. 5d-f). The analysis aimed to explore the biological processes and functions associated with these computationally selected proteins. The enrichment results showed that these predicted substrates were associated with synaptic function, neuronal signal transduction, and transmembrane molecular exchange (Fig.5d). These associations are consistent with previous studies reporting roles of Caspase-3 in the nervous system(54–56). The predicted substrates also exhibited functional profiles closely aligned with those of the experimentally validated substrates (Fig.5e). Both sets exhibited enrichment in terms such as trans-synaptic signalling (GO:0099537), synaptic signalling (GO:0099536), synaptic membrane (GO:0097060), glutamatergic synapse (GO:0098978), alpha-tubulin binding (GO:0043014), and tubulin binding (GO:0015631). These shared enrichment patterns indicate that the predicted candidates and experimentally validated substrates are associated with several similar biological functions. However, functional enrichment does not independently confirm that the candidate proteins are cleaved by Caspase-3 at the predicted sites. In addition, the predicted substrates were enriched for terms related to organic cation transport, amino acid transmembrane transporter activity, and organic acid transport. These results suggest additional biological processes that may be associated with the predicted substrate set. They provide possible directions for further investigation but do not establish new functions of Caspase-3.

### Experimental validation confirms three new Caspase-3 substrates that were predicted by OmniCleave

We selected a few previously unannotated candidate caspase-3 substrates generated by OmniCleave, including CUL7, THOC5, and RPIA, for validation using in vitro cleavage assays (Fig. 6a, Supplementary Table S8, and Supplementary Material). MS/MS Spectrum plots are shown in Supplementary Figures S1-S30. The unannotated putative substrate proteins were incubated either alone (control) or with Caspase-3 (test). SDS-PAGE analysis revealed a marked reduction in the intensity of all three substrate bands in the test reactions compared to the controls, confirming proteolytic cleavage (Fig. 6b and 6f). Subsequent LC-MS/MS analysis identified 12, 6, and 12 cleavage sites for THOC5, RPIA, and CUL7, respectively. Of these, OmniCleave correctly predicted 8, 5, and 8 sites. To compare, the recently published structure-based Procleave predicted only 0, 2, and 2 sites, respectively. SeqLogo analysis showed that the identified cleavage motifs (THOC5: VLSD; RPIA: DEID; CUL7: SLLD) are consistent with Caspase-3’s preference for acidic residues (Fig. 6c).

**Figure 6.**
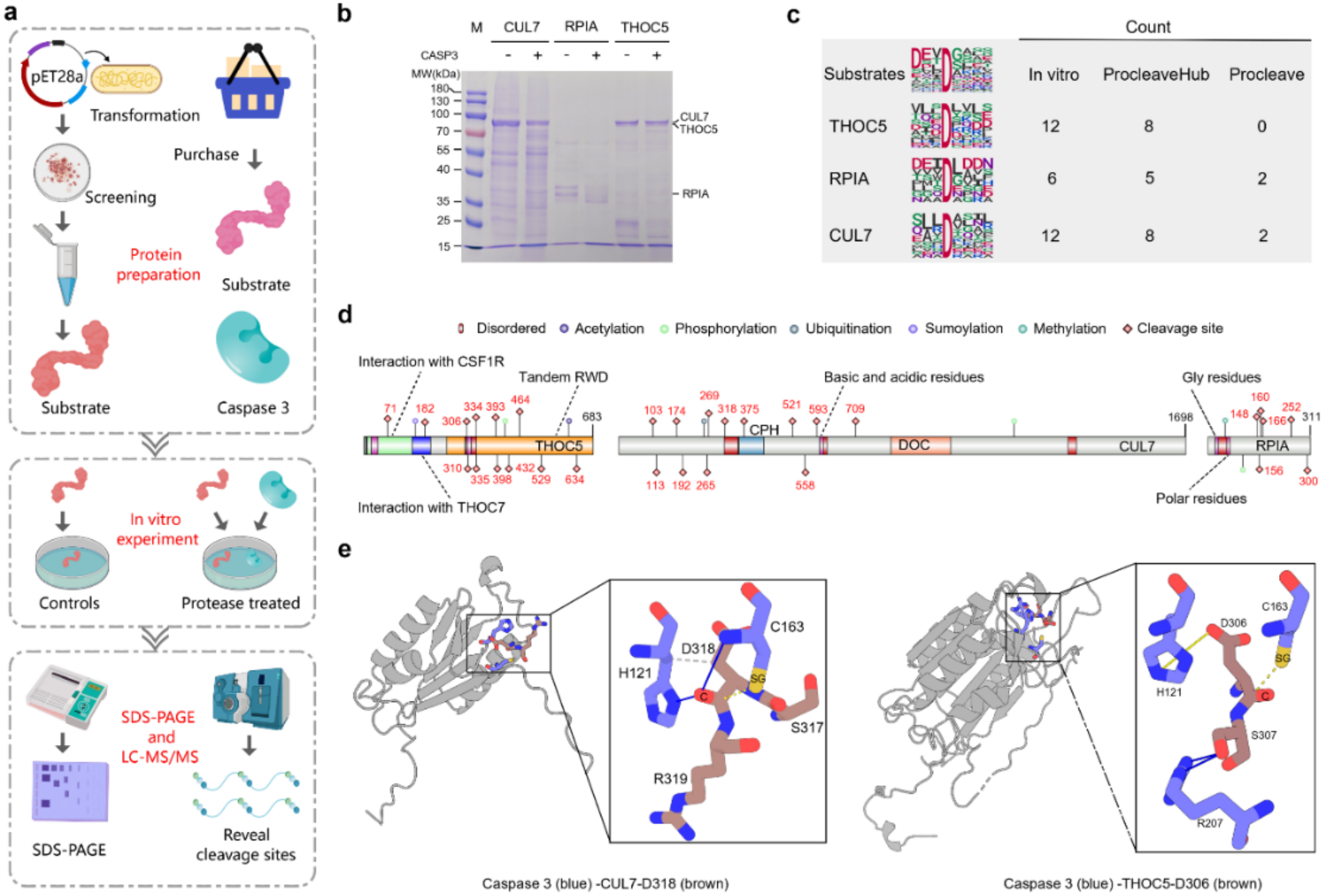
*In vitro* cleavage assays validate OmniCleave predictions. **a.** Experimental workflow: substrate proteins were prepared via E. coli expression or commercial purification, followed by in vitro cleavage assays (control vs. experimental), SDS-PAGE, and LC-MS/MS analysis. **b**. SDS-PAGE results show markedly reduced bands for THOC5, RPIA, and CUL7 in experimental groups, indicating cleavage. **c**. Cleavage sites detected in vitro are compared with predictions from OmniCleave and Procleave; Seqlogo plots illustrate P4-P1 sequence preferences of validated substrates and three candidate proteins. **d**. Cleavage site distribution across protein domains: THOC5 sites localise mainly to the Tandem RWD domain, while CUL7 and RPIA sites are predominantly in non-domain regions. **e**. Exploratory visualization of putative Caspase-3-substrate complex structures. Representative constraint-guided HelixFold3 model of the Caspase-3-THOC5 complex and GRAMM docking model of the Caspase-3-CUL7 complex. The models illustrate the spatial arrangements of Caspase-3 residues H121 and C163 relative to the predicted THOC5 D306 and CUL7 D318 cleavage regions. Putative intermolecular contacts were identified using PLIP and visualized using PyMOL. Both models were generated using predefined residue constraints and are presented as exploratory, hypothesis-generating structures. The predicted contacts represent possible interactions in the selected models but do not establish native binding modes, the structural basis of substrate recognition, or catalytic mechanisms.

Domain annotation (from the UniProt database) of Caspase-3 substrates revealed that cleavage sites of THOC5 predominantly reside within the Tandem RWD domain, including regions interacting with CSF1R and THOC7, whereas CUL7 and RPIA cleavage sites were largely located in non-domain regions (Fig. 6d). These observations indicate that substrate specificity is shaped not only by canonical sequence motifs but also by local structural context and flexible regions. To explore possible spatial arrangements of the predicted cleavage regions, we generated exploratory complex models for Caspase-3 with CUL7 and THOC5 (Fig. 6e). These models were obtained using constraint-guided complex prediction and protein–protein docking, followed by interaction analysis and structural visualization. For the Caspase-3-CUL7 complex, the selected GRAMM docking pose positioned the predicted CUL7 D318 cleavage region near the catalytic residues C163 and H121 of Caspase-3(57, 58). In this model, C163 was located close to the predicted cleavage region, while H121 was positioned near the substrate. For the Caspase-3-THOC5 complex, the selected HelixFold3 model placed the predicted THOC5 D306 cleavage region close to C163, with H121 positioned near the substrate and a putative hydrogen bond identified between Caspase-3 R207 and THOC5 S307. Putative hydrogen bonds, salt bridges, and hydrophobic contacts in the selected models were identified using the Protein-Ligand Interaction Profiler (PLIP)(59) and visualized in PyMOL(60). As both models were generated using predefined residue constraints, the predicted arrangements may be influenced by these constraints. We therefore present these structures as exploratory, hypothesis-generating models rather than evidence of native binding modes.

## Discussion

OmniCleave is an innovative and comprehensive computational framework for parsing protease-substrate interactions and identifying substrate cleavage sites. By integrating protease-protease interaction networks with advanced protein language model embeddings and hierarchical subgraph representations, OmniCleave provides a comprehensive view of proteolytic regulation that surpasses the capabilities of existing methods, encompassing both structure and sequence. Unlike traditional sequence-based approaches, OmniCleave dynamically captures the determinants of proteolytic specificity at the structural level, achieving precise predictive capability and interpretability. Integrating 3D structural information at different modalities, including atomic and residue levels, enhances the biological relevance of cleavage predictions. Local structural context modelling enables OmniCleave to account for cleavage-relevant conformations, residue proximity, and interaction geometry, while avoiding the computational overhead and redundant information associated with modelling the entire protein structure. These factors are often overlooked in previous sequence-based approaches. Furthermore, by modelling protease-protease interaction networks, OmniCleave effectively simulates protease co-cleavage patterns. This modelling approach enables OmniCleave to overcome key limitations of existing methods, achieving single-pass inference for substrate cleavage site identification across 103 proteases and protease-substrate interaction recognition. The ablation results demonstrate that local P4-P4’ sequence specificity is a major source of predictive information for protease cleavage-site prediction, which is biologically expected given the strong residue preferences of many proteases around the scissile bond. The Seq + PPI variant retained strong predictive performance, confirming that local sequence information provides a substantial fraction of the predictive signal; however, its performance remained clearly below that of the complete OmniCleave model. Conversely, removing the sequence-derived features caused the largest performance reduction, while the remaining structural, energetic, and protease-relation features still retained predictive information. In addition, removal of the graph-based structural module, DSSP-based features, or energy-based features consistently reduced performance relative to the complete model. Together, these complementary ablations indicate that local sequence specificity provides a major predictive foundation, whereas graph-based structural representations, secondary-structure descriptors, energetic features, and protease-context information provide additional discriminatory information beyond the local sequence window. Mechanism analysis demonstrates that OmniCleave accurately captures the effect of distal residues on cleavage specificity. Feature perturbation experiments further demonstrate the complementary roles of energy-related and structural descriptors. Crucially, the model successfully reproduces experimentally validated protease-substrate relationships and uncovers previously uncharacterised substrate functions, demonstrating its ability to generate biologically meaningful hypotheses. Validation using in vitro caspase-3 cleavage experiments confirmed OmniCleave can identify previously unannotated cleavage events.

Despite OmniCleave’s strong performance, several limitations warrant attention. The current framework relies on the accuracy of predicted protein structures, the completeness of existing protease-substrate and protein-protein interaction datasets, and the scale of curated hydrolysis data. OmniCleave uses structural features derived primarily from AlphaFoldDB and ESMFold models. Although the pLDDT-stratified evaluation showed that the model retained predictive performance across different confidence levels, predicted structures in flexible or low-confidence regions may not accurately represent the conformational states accessible in vivo. Moreover, pLDDT reflects model confidence rather than a direct experimental measurement of structural disorder. Future work incorporating experimentally determined structures, conformational ensembles, or disorder-aware representations may further improve predictions in structurally dynamic regions, thereby characterising the true conformation of protease-substrate complexes and further enhancing the explanatory power of proteolysis mechanisms from a structural perspective.

Rosetta energy-based features were calculated directly from unrelaxed AlphaFold DB and ESMFold structures to maintain the computational scalability of OmniCleave. These descriptors may therefore reflect not only the local physicochemical environment but also geometric imperfections or side-chain clashes in the predicted structures. Accordingly, they should not be interpreted as relaxed-state energies or physical free-energy estimates. Nevertheless, the no-energy ablation results indicate that the overall performance of OmniCleave is not exclusively dependent on these descriptors. Additionally, computational demands associated with structural feature extraction could pose challenges for large-scale or high-throughput applications. The contribution of the PPI module may depend on the coverage and quality of available protease interaction annotations. Proteases with limited PPI information may benefit less from relational information than well-characterised proteases. Nevertheless, OmniCleave can still make predictions using protease- and substrate-derived sequence and structural features, even when PPI annotations are incomplete. More complete and context-specific interaction networks may further improve prediction performance for poorly characterised proteases.

In summary, OmniCleave establishes a unified platform (Screenshots of the webserver and GUI are shown in Supplementary Figure S31) to advance research on protease hydrolysis mechanisms, substrate function annotation, and protease synergistic mechanisms. By bridging structural, physicochemical properties, and protease-protease information within a single architecture, it provides an effective pathway for data-driven discovery in proteolytic biology and offers actionable hypotheses for biological experiments. Moreover, its ability to perform structural-level dissection of protease-substrate interactions enables the reasonable validation of the design effectiveness of multi-targeted small-molecule inhibitors, optimises protease-substrate design, and facilitates the de novo design of proteases with customised catalytic properties. This method can link computational predictions with experimental protease biology.

## Materials and Methods

### Protease-substrate cleavage dataset construction

This study utilized proteolytic cleavage events curated in the MEROPS database (Release 12.4)(39). Each record contained a protease identifier, a substrate UniProt accession, a protease Uniprot accession, and an annotated cleavage position (the position of the P1 residue). To ensure that we have a sufficient amount of data for proper training and testing, we excluded proteases with fewer than 30 cleavage sites. We also used CD-HIT(61) to reduce sequence similarity among substrate proteins. For the standard dataset, substrate sequences were clustered using a 70% sequence identity threshold, and one representative protein was retained from each cluster. After filtering, we collected 57,278 protease-cleavage site pairs, involving 103 proteases and 9,651 substrates. The tertiary structures of proteases were obtained from the AlphaFold Protein Structure Database (AlphaFold DB)(41, 42). As for the substrates, 9,226 tertiary structures were retrieved from AlphaFold DB, while the remaining 425 were generated using ESMFold(43) to complete the dataset.

For each substrate protein, we first collected all positions annotated as cleavage sites for any of the 103 proteases. Thes position were treated as positive sites. They were excluded from the candidate negative pool, regardless of which protease was associated with the annotation. A position could be selected as a putative negative site only if it had no cleavage annotation for any of the 103 proteases. For each substrate protein, the number of sampled putative negative sites was matched to the number of annotated cleavage sites on that protein, thereby maintaining a 1:1 positive-to-negative ratio at the substrate-protein level. Negative sites were randomly sampled without replacement from the candidate negative pool, while ensuring that no annotated cleavage site was included as a negative example. Thus, 57,278 putative negative sites were randomly sampled in total, maintaining a 1:1 positive-to-negative ratio both at the substrate level and across the complete dataset.

After sample constructin, the dataset was divided into training and independent test sets at the substrate-protein level. All positive and negative samples from the same substrate protein were assigned to the same partition. Consequently, no substrate protein was shared between the training and test sets. We also verified that no unique cleavage-site record, defined by the combination of substrate UniProt accession and cleavage position, occurred in both sets. Because only one representative protein was retained from each CD-HIT cluster, highly similar substrate proteins with more than 70% (30%) sequence identity were also prevented from being distributed across the training and test sets. The final dataset was divided into training and independent test sets at a ratio of 7:3, resulting in 80,168 and 34,388 samples, respectively. Subsequently, 10% of the training set was selected using stratified sampling to construct a validation set for model optimisation during training. In addition to this redundancy-reduced dataset, we constructed a more stringent data partition using a 30% sequence-identity threshold to evaluate model generalization.

We also examined whether model performance was affected by the negative-sampling strategy. In addition to the original random-negative dataset, we constructed two alternative datasets using far-site and near-site negative sampling. In the random-negative setting, negative sites were selected from all eligible unannotated positions without restrictions on their distance from known cleavage sites. For near-site negative sampling, candidate negative positions were selected from unannotated residues located 1-8 residues away from an annotated cleavage site on the same substrate. These sites were close to known cleavage positions and could share similar local sequence and structural environments. For far-site negative sampling, candidate negative positions were required to be at least 15 residues away from every annotated cleavage site on the same substrate. In both cases, distance was measured along the amino-acid sequence rather than in three-dimensional space. All three negative-sampling strategies followed the same basic rules. Positions annotated as cleavage sites for any of the 103 proteases were excluded from the negative pool. The number of sampled negatives for each substrate protein equalled the number of annotated cleavage sites on that protein, ensuring that the positive-to-negative ratio was identical across the three sampling schemes. Only positions with complete P4-P4’ sequence windows were considered. The same positive samples and substrate-level partitioning rules were used across the three strategies. Thus, the main difference among the random-, far-site-, and near-site-negative datasets was the rule used to select candidate negative positions. For both the far-site and near-site strategies, five independent negative-sampling runs were performed using different random seeds.

### Neural network of OmniCleave

In this section, we detail the modules of OmniCleave. OmniCleave comprises three modules: a cleavage-centric hierarchical encoder, a protease-protease interaction (PPI) module, and a graph transformer-based protease-substrate interaction module. For the cleavage-centric hierarchical encoder, we use an equivariant graph neural network to learn the local three-dimensional structural information of the cleavage site. For the protease-protease interaction (PPI) module and the Graph transformer-based protease-substrate interaction module, we introduce protease interaction information (PPI) and use a heterogeneous graph representation to model the interaction between proteases and substrates (cleavage site), respectively. We convert the prediction task of the cleavage site into a link prediction task between the protease and substrate cleavage sites, realising the prediction capability of the relationship between multiple proteases and one site.

### Cleavage-centric hierarchical encoder

To capture structural and functional characteristics of proteolytic cleavage, we employed a fixed-radius approach to construct local structural subgraphs representations centered on cleavage sites for samples in the training, validation, and test sets, encompassing both residue-level and atom-level representations. Message passing occurs only within the respective subgraphs. Compared to modelling the entire substrate structure, this localised representation not only reduces complexity and computational cost but also retains critical contextual information relevant to cleavage activity. Our constructed subgraphs offer a more detailed representation of the surrounding context by integrating sequence, geometric, and energetic information. In detail, for a given substrate cleavage site, we extracted residues whose alpha carbon (Cα) atoms are within a 10 Å Euclidean distance from the Cα atom of the cleavage site residue (P1). The resulting local environment was modelled as a hierarchical subgraph *G*^*C*^, incorporating two levels of granularity: a residue-level subgraph *G*^*R*^ and an atom-level subgraph *G*^*A*^. Formally, this can be expressed as:

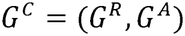

### Residue-level subgraph

The residue-level subgraph *G*^*R*^ =(*V*^*R*^,*E*^*R*^), consists of nodes *V*^*R*^ and edges *E*^*R*^. Specifically, a residue i was regarded as a node *v*_*i*_ ∈ *V*^*R*^ and was represented by three types of features: (i) sequence embeddings obtained from the ESM-2 protein language model (ESM2-650M)(43), encoding context-aware residue representations; (ii) geometric features based on DSSP annotations(62), describing features such as a α-helices, β-strands, solvent exposure, and others (summarised in Supplementary Table S5); and (iii) energy terms computed via PyRosetta 4 (release 2024.24) with the default all-atom score function, including van der Waals, solvation, electrostatic, backbone, and side-chain conformational terms(49). Although the Rosetta energy terms are calculated from atomistic coordinates, we did not manually aggregate atomic energies. Instead, we used the native residue-level score decomposition for each residue returned by the function provided in PyRosetta. We extracted 12 Rosetta score components: *fa_intra_sol_xover4, fa_intra_rep, rama_prepro, omega, p_aa_pp, fa_dun, ref, fa_atr, fa_rep, fa_sol, fa_elec*, and *lk_ball_wtd* (see Supplementary Table S6 for details). We scored the predicted structures directly without structural relaxation, and used these values as conformation-dependent energetic descriptors rather than estimates of thermodynamic stability or binding free energy. Collectively, these features define a residue representation as:

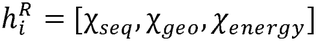

An edge (*i,j*) ∈ *E*^*R*^ between residues *i*and *j* was formed if their *Cα* atoms were within an 8Å Euclidean distance, with a maximum of 10 edges per residue. Each edge was represented by a radial basis function (RBF) distance expansion and a unit direction vector:

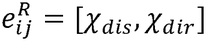

### Atom-level subgraph

The atom-level subgraph *G*^*A*^ =(*V*^*A*^,*E*^*A*^), refines the residue-level subgraph *G*^*R*^ by decomposing each residue into its constituent atoms. In this subgraph, each atom m was regarded as a node, and was represented by learnable embeddings that encode its atomic type (e.g., carbon, oxygen, nitrogen) and its position distribution within the residue:

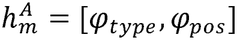

Edges in *G*^*A*^ follow a similar construction to *G*^*R*^, connecting atoms within 8 Å, with at most 10 neighbours per atom. The edge representation between atom m and atom n was defined as:

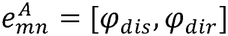

### Multi-Scale Subgraph Update for Cleavage Sites

We utilize the Generalist equivariant transformer GET(63) to update hierarchical cleavage site subgraphs by capturing information at both residue and atom-level. Given a residue i with its initial feature representation 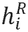, the residue contains *S*^(i)^ atoms with the atom features denoted as:

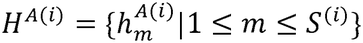

We first apply GET to refine atom features:

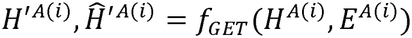

where *H*′^*A*(*i*)^ represents the updated atom feature set, and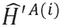serves as its global representation (atom-level graph features), obtained by summing over *H*′^*A*(*i*)^; *E*′^*A*(*i*)^ represents the atom edges with their attributes. Then, we aggregate the atom features to obtain the feature of residue *i*:

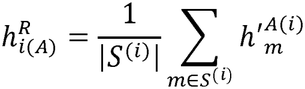

where 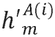 denotes the updated feature of the m-th atom. The resulting residue feature 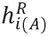 is then concatenated with the initial residue feature 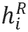:

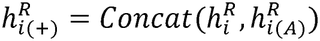

After atom-level updating, another GET layer is applied to further update the residue features, where the residue feature set are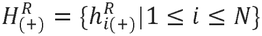, and *N* is the number of residues in the subgraph *G*^*C*^:

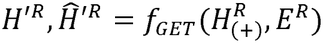

where *H*′^*R*^ represents the updated residue feature set, and 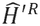 serves as the global representation (residue-level graph features), which can be interpreted as a compressed representation of subgraph *G*^*C*^; *E*^*R*^ represents the residue edges with their attributes.

### Generalist equivariant transformer (GET)

GET (63) was employed as a core component to update the cleavage subgraph *G*^*C*^. GET introduces a bilevel unified representation to model molecular complexes as geometric graphs of sets. This design allows the processing of variable-size blocks, overcoming the limitations of conventional single-level representations or hierarchical pooling methods based on fixed-size vector-form features. The unified representation of GET accommodates matrix-form node features and 3D coordinates while ensuring E(3)-equivariance and permutation invariance within the block. GET consists of three E(3) equivariant components: a bilevel attention module, a feed-forward module, and an equivariant layer normalization module. The bilevel attention module captures both sparse interactions at the block-level (e.g., residues) and dense interactions at the atom-level. The feed-forward module updates the hidden states and coordinates of each atom, and the equivariant layer normalization module normalizes the hidden states and coordinates.

Specifically, a cleavage site subgraph *G*^*C*^ that contains blocks (residues) can be defined as a geometric graph of sets:

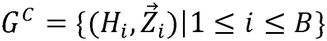

where *H*_i_ represents a set of atom feature vectors and 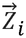 denotes the 3D atom coordinates. First, the bilevel attention module updated the hidden state and coordinates at the atom level and residue level. The **atom-level cross-attention** *α*_*ij*_ from residue *j* (contains *n*_*j*_ atoms) to residue *i* (contains *n*_*i*_ atoms) was calculated as follows:

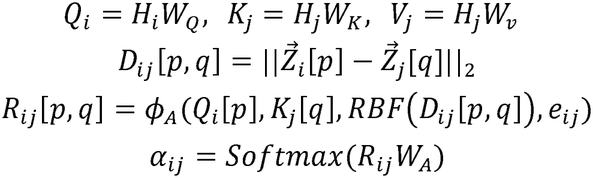

where *W*_*Q*_, *W*_*K*_, *W*_*v*_,*W*_*A*_ are trainable matrices;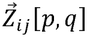, represents relative coordinates between any two atoms *p* (in residue *i*) and *q* (in residue *j*),*D*_*ij*_ [*p,q*] is their distance; *e*_ij_ denotes edge features; *ϕ* _*A*_is a multi-layer perceptron (MLP); and RBF denotes radial basis functions embedding. Then the **block-level cross-attention** *β*_*ij*_ from *j* to *i* was calculated as:

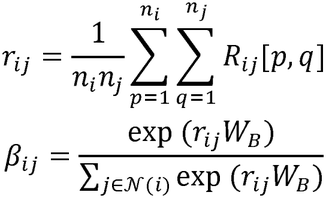

where *W*_*B*_ is a learnable matrix and *N*(*i*) denotes neighborhood residues of *i. r*_*ij*_ represents the global relation between *i* and *j*. With the computed attention values *α*_*ij*_ and *β*_*ij*_, the hidden features and coordinates of atom in residue *i* were updated as:

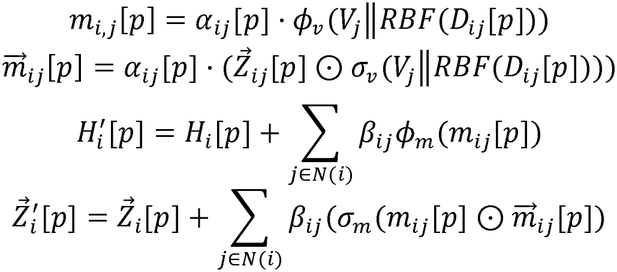

where ∥ denotes concatenation along the second dimension; *ϕ*_*v*_, *σ*_*v*_, *ϕ*_*m*_, *σ*_*m*_ are MLPs; and ⊙ denotes the element-wise multiplication. The hidden features of residue are updated in same way, except that the average of the atomic hidden states (the average of all atomic hidden states corresponding to the residue) is used as the supplement to the initial state (initial feature) of the residue. The coordinates of the residue are set to the Cα atom coordinates.

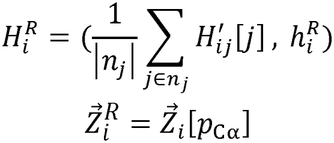

where 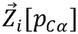 denotes the coordinates of the Cα atom in residue *i*.

In the **feed-forward layer**, the *H*_*i*_ and 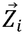 for each atom are updated individually:

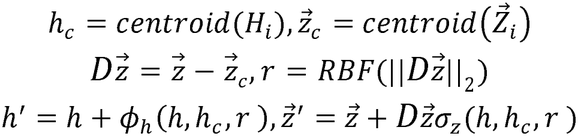

where the *h*_*c*_ and 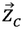 denotes the centroids of the block; 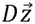 represents the relative coordinate between each atom and the centroid; *r* is the representation of distance; *ϕ*_*h*_ and *σ*_*z*_ are MLPs. The **equivariant layer normalization layer** not only considers E(3)-equivariance but also stabilizes and accelerates the training of deep neural networks. The hidden vectors and the coordinates of individual atoms are normalized as follows:

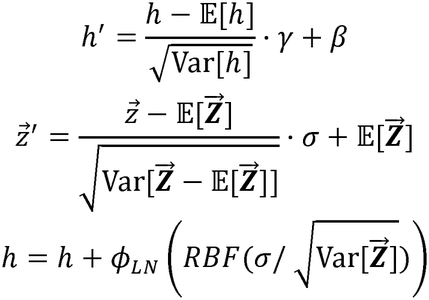

where 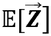 denotes the centroid of the entire graph; 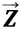 represents the coordinates of all atoms in all blocks (residues); *γ, β*, and σ are learnable parameters; *ϕ*_*LN*_ is an MLP, and the 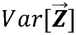 computes the change in all atomic coordinates relative to the center of mass.

### Protease-protease interaction (PPI) module

Structural and catalytic similarities among proteases often lead to recognition and cleavage of overlapping substrate sites. To incorporate these functional associations, we modelled a graph *G*^*P*^ =(*V*^*P*^,*E*^*P*^ ) of protease-protease interactions (PPI) as one of the model inputs. Proteases were regarded as nodes, and protease interactions were denoted as unweighted edges. The representation of each node was computed by averaging the residue-level features across residues within its peptidase unit(39). For a given protease *T*_*d*_ (1 ≤ *d* ≤ 103), its feature vector 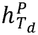 can be formulated as:

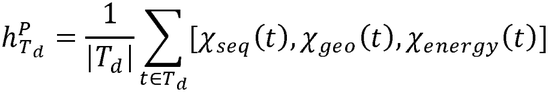

where denotes a residue within *T*_*d*_, and | *T*_*d*_ | is the total number of residues within the peptidase unit. The terms *X*_*seq*_ (*t*), *X*_geo_ (*t*), and *X*_*energy*_ (*t*) correspond to ESM-2 sequence embeddings, DSSP-derived geometric features, and Rosetta energy features, respectively.

### Graph transformer based protease-substrate interaction module

#### Interaction modeling between proteases and substrates

To characterize the intrinsic mechanism of proteolytic catalysis, we established connections between proteases and substrate cleavage sites. Specifically, the protease feature set *H*^*P*^, corresponding to a node set *V*^*P*^ (contains 103 proteases), is first transformed via an MLP layer:

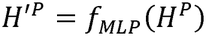

Next, the protease nodes *V*′^*P*^are linked to 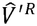, where *V*′^*P*^ contains the updated feature set *H*′^*P*^, and 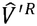 corresponds to the node set of 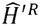:

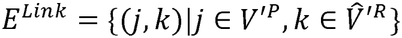

This process can be regarded as constructing a new graph by integrating protease-substrate interactions (protease-cleavage site interaction graph), denoted as *G*^*P R* =^(*V*^*PR*^,*E*^*PR*^), where ^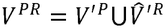^and *E*^*PR*^*=E*^*p*^ U*E*^*Link*^. The *G*^*PR*^ is then processed by a two-layer Graph Transformer Convolutions (GTC)60 for message passing:

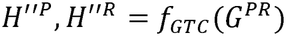

After this step, the updated features of *H*”^*P*^ and *H*”^*P*^ will be used to calculate the probability of whether there is a link between the protease and the substrate cleavage site.

### Relation prediction between PPI and cleavage sites

We integrate multimodal information from proteases and substrates to estimate the probability of cleavage. For proteases, we consider an embedded representation of the protein language model of proteases. For substrate proteins, in addition to structural features, we also considered the sequence-based features (chemical group feature C^R^ and the amino acid (AA) type feature S^R^)(27). Given the potential cleavage sites, a sliding window approach (P4-P4’) is used to extract the local sequence features, and the chemical groupings of AA are used to calculate chemical features. These two features are coded as embedded according to the type and chemical grouping of amino acids, respectively. The probability of the relationship *Pro*^*PR*^ is calculated as follows:

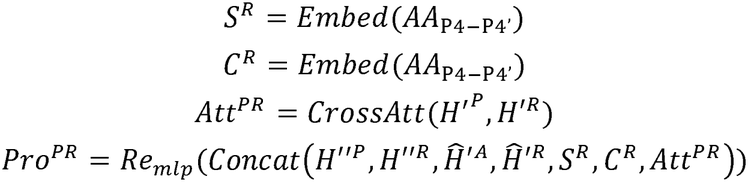

where *AA*_P4-P4’_ represents the P4-P4’ local sequence surrounding the potential cleavage sites; *H*^“P(i)^ denotes the protease i; A*tt*^*PR*^ represents cross-attention of potential cleavage sites to all proteases;*Re* _*mlp*_ is an MLP.

### OmniCleave hyperparameters and training

Hyperparameter tuning for OmniCleave was conducted experientially through iterative exploration over the training dataset. We use GET as the basis for the graph neural network. In our experiments, we used a GET layer in the graph encoder, a hierarchical graph setting, and 128 hidden layer sites, and the output dimension is 256. For proteases, the extracted language model features, energy features, and structural features are reduced to 256 dimensions through a double-layer fully connected layer, which is used to initialize the protease-cleavage site interaction graph. Finally, OmniCleave uses a fully connected layer with 1047 input dimensions to receive multi-source fusion features to identify the cleavage site of the protease.

OmniCleave was trained on PyTorch 1.13 and the parameters were initialized using Xavier initialization(64). To prevent overfitting, in addition to LayerNorm in the GET module, we also apply dropout (0.2) between linear layers. The network was trained using an SGD optimizer with an initial learning rate of 0.0003 and a weight decay of 1e-3. The model was trained with a batch size of 64 for 200 epochs using the cross-entropy loss function. To evaluate the model, we utilized a range of evaluation metrics commonly employed in bioinformatics, including the F1 score, AUC, and AUPR. Additionally, we employed the learning rate adjustment strategy (ReduceLROnPlateau) to address the optimisation challenges encountered during model training. Our model was trained on an NVIDIA GeForce RTX 3090 GPU equipped with 24 GB RAM.

### Construction of the protease-protease interaction network

Protease-protease interaction information was obtained from the STRING database(44) (version 11.5). The proteases included in the OmniCleave dataset were mapped to their corresponding STRING protein entries using gene names. We collected interaction records supported by knowledge-based evidence, assay-based predictions, and genomic predictions. Interactions were selected using the STRING confidence score, with a minimum threshold of 0.400. Only interactions meeting this threshold were retained to construct the protease– protease interaction network.

Each protease was represented as a node, and the retained interactions were represented as unweighted edges. Proteases without qualifying STRING interactions were kept in the graph but had no protease–protease edges. These proteases were connected only to cleavage-site nodes through edges defined by the annotated protease–cleavage-site relationships. The resulting graph was used to provide protease interaction information for OmniCleave. As STRING includes functional associations from different evidence sources, its interactions do not necessarily represent direct physical binding between proteases.

### Comparison with AlphaFold

AlphaFold3 was proposed for predicting the structure of proteins in complex with one or more other proteins, nuclei, or small molecules. In this study, we used AlphaFold3 to perform cleavage site identification experiments based on the assumption that the cleavage site lies within the binding region of the protease and its substrate protein. First, we used AlphaFold3 to predict the structure of the protease-substrate complex using the sequences of the protease and substrate. Subsequently, the optimal complex structure was selected to assess whether AlphaFold3 correctly predicted the substrate’s cleavage site. If the known cleavage site (P1) is located within the complex binding region (judged using PyMol) and its minimum spatial distance to the amino acids in the protease (not limited to the enzyme active site) is less than 5 angstroms, then we consider AlphaFold 3 to have correctly predicted the site.

### pLDDT grouping analysis

To assess the influence of predicted-structure confidence, the per-residue pLDDT value at the P1 position was extracted for each cleavage and non-cleavage sample. Test samples were divided into low-confidence (pLDDT < 70), medium-confidence (70 ≤ pLDDT < 90), and high-confidence (pLDDT ≥ 90) groups. Model performance was evaluated independently in each group using AUC and MCC. Distributional analyses included all available samples, whereas the performance-stratification analysis was restricted to the independent test set.

### Ablation analysis

To systematically evaluate the contributions of the different model components and information sources, we compared the complete OmniCleave model with six ablation variants. The single-level variant retained the residue-level graph representation but removed the atom-level subgraph and the corresponding hierarchical integration between the residue and atomic levels. The w/o PPI variant removed the PPI-based protease interaction module while preserving all substrate-derived sequence and structural representations. In the w/o GNN variant, the GET encoders applied to the local residue-level and atom-level substrate graphs were removed. The P4-P4’ sequence-derived representations, DSSP-based features, energy-based features, and PPI information were retained in this variant. Thus, the w/o GNN setting specifically assessed the contribution of graph-based encoding of the local structural environment. The w/o DSSP variant removed the DSSP-based features while retaining all other model inputs. The w/o energy variant removed the energy-based features. Finally, the w/o seq_based variant removed the P4-P4’ amino-acid and chemical-group representations, denoted as S^R^ and C^R^, while retaining the graph-based structural, secondary-structure, energetic, and protease-relation features. In addition, we constructed a Seq + PPI variant to evaluate the predictive information provided by the local P4-P4’ sequence window in the absence of substrate structural information. In this setting, only the P4-P4’ amino-acid type representation SR and chemical-group representation CR were retained on the substrate side, while the residue- and atom-level GET/GNN representations, DSSP-derived descriptors, and Rosetta-derived score components were removed. The PPI-based protease representation was retained to preserve protease-specific conditioning within the unified 103-protease model. Without protease-specific information, identical substrate sequence windows associated with different proteases would be indistinguishable to the model. The Seq + PPI variant therefore evaluates sequence-based substrate recognition while maintaining the protease context required for multi-protease prediction. All ablation variants were trained and evaluated using the same data partitions, optimisation settings, and evaluation procedures as the complete OmniCleave model. Each experiment was repeated over five independent runs, and performance was assessed using AUC, AUPR, and F1 score.

### Motif-overlap analysis

To assess whether exact local sequence motifs were shared between the training and test sets, each valid sample was represented by its complete eight-residue P4-P4’ sequence window. Exact motif overlap was defined as an identical P4-P4’ amino-acid sequence occurring in both partitions. We first calculated the numbers of unique P4-P4’ motifs in the complete training and test sets, irrespective of sample label, and determined the size of their intersection. We subsequently repeated the analysis for positive samples only. A shared positive motif was defined as an identical P4-P4’ sequence occurring with label 1 in both the training and test sets.

### Feature Importance

To effectively determine the importance of specific node features, we refer to ref(29), perturb the value of each node feature in the corresponding protease test sample, and examine how much the test loss value decreases, that is, the absolute value of (original loss - perturbation loss). Through the above steps, we can evaluate the importance of all node features.

### Statistical test

For each putative negative-sampling strategy, five independently sampled datasets were generated, and OmniCleave was retrained and evaluated separately on each dataset. The analysis included seven representative proteases (caspase-3, caspase-7, MMP-2, MMP-9, MMP-7, granzyme B, and thrombin) and three evaluation metrics: AUC, AUPR, and F1 score. Because several comparison tools support only a subset of these proteases, each statistical comparison was restricted to the proteases supported by both OmniCleave and the corresponding tool. Unsupported proteases were not treated as missing performance values or assigned artificial scores. For each supported protease and independent replicate, we calculated the paired performance difference as:

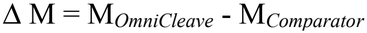

where M denotes AUC, AUPR, or F1 score. A positive value therefore indicates better performance by OmniCleave. Statistical significance was assessed using a one-sided Wilcoxon signed-rank test, with the alternative hypothesis that OmniCleave achieved higher performance than the corresponding comparison method. The resulting P-values were adjusted for multiple comparisons using the Benjamini-Hochberg procedure, and adjusted q-values below 0.05 were considered statistically significant.

### Visualization of putative Caspase-3-substrate interactions

For the Caspase-3-THOC5 complex, constraint-guided complex prediction was performed using HelixFold3(65) with 10 recycling iterations. An approximately 5 Å distance constraint was imposed between Caspase-3 C163 and the predicted THOC5 cleavage residue D306. Five candidate complex models were generated, and the top-ranked model satisfying the imposed constraint was selected for subsequent structural visualization. For the Caspase-3-CUL7 complex, protein-protein docking was performed using the GRAMM Web Server(66). The Caspase-3 receptor structure was obtained from the Protein Data Bank (PDB ID: 2XYG), whereas the CUL7 ligand structure was predicted using AlphaFold3. Caspase-3 residues C163 and H121 and CUL7 residue D318 were specified as interface residues. Ten docking poses were generated, and the top-ranked pose compatible with the defined interface was selected for subsequent analysis. Putative non-covalent contacts in the selected complex models were identified using the online Protein-Ligand Interaction Profiler (PLIP-2.2.0)(59), including predicted hydrogen-bonding, salt-bridge, and hydrophobic interactions. All interactions were identified using the default geometric thresholds provided by the PLIP web server, without manual modification. Specifically, hydrogen bonds were identified using a maximum donor-acceptor distance of 4.1 Å and a minimum donor-hydrogen-acceptor angle of 100°. Salt bridges were identified between oppositely charged groups when the distance between their charge centers was within 5.5 Å. Hydrophobic contacts were detected between hydrophobic carbon atoms within a distance of 4.0 Å. The resulting complex structures and interaction annotations were visualized using PyMol(60). These analyses were performed to illustrate plausible interaction geometries around the predicted cleavage regions and were not used to establish native binding modes, binding affinities, or catalytic mechanisms.

### Proteome-wide prediction of human protease substrates

To identify previously unannotated human protease substrates, we collected human protein structures from the AlphaFold Protein Structure Database. Proteins with annotated cleavage sites in MEROPS (Release 12.4) were excluded, regardless of the associated protease. We also removed proteins without available structural models.

After filtering, 14,332 human proteins were retained for prediction. We applied OmniCleave to these proteins using the human proteases included in our dataset. For each protein, the model calculated cleavage probabilities at all eligible positions for each of the human proteases. Only positions with a complete P4-P4’ sequence window were considered. A protein was selected as a candidate substrate if at least one position had a predicted cleavage probability greater than or equal to 0.99 for any of the human proteases under consideration. Using this criterion, we identified 1,379 unique human proteins. Functional enrichment analysis was performed on these 1,379 proteins using the Metascape website(46).

### Enrichment analysis

We used the Metascape website(46) to perform pathway and Gene Ontology enrichment analysis on the predicted substrate proteins, with the parameters set as “Min Overlap = 3, Min Enrichment =1.5, pvalueCutoff = 0.01, qvalueCutoff = 0.05”. We used the default background provided by Metascape without specifying a custom background gene set.

### In vitro cleavage assays validation

#### Protein preparation

Full-length recombinant human Caspase-3 (CASP3, UniprotKB: P42574) and RPIA (UniprotKB: P49247) used in this study were purchased commercially (CASP3: MedChemExpress #HY-P701341; RPIA: FineTest #P5260). Both CASP3 and RPIA were expressed and purified in *Escherichia coli* (*E*.*coli*), and were supplied with greater than 90% purity as determined by SDS-PAGE. CASP3 was provided as a 0.22 μm-filtered solution in buffer containing 50 mM Tris-HCl (pH 7.5), 200 mM NaCl, 20% glycerol, and 1 mM DTT. The enzymatic activity of Caspase-3 was validated by its ability to cleave the synthetic substrates Ac-DEVD-AMC (specific activity: 26.57 U/mg) and Ac-DEVD-AFC (specific activity: 29,332.2 pmol/min/μg) under defined assay conditions as claimed by the manufacturer. RPIA was supplied in lyophilised form from a 0.2 μm-filtered solution containing 10 mM HEPES, 150 mM NaCl, and 5% trehalose, at pH 7.4. Prior to use, the lyophilised protein was reconstituted in Tris-HCl buffer (20 mM Tris-HCl, pH 8.0, 500 mM NaCl). Full-length THOC5 (UniprotKB: Q13769) and a truncated form of CUL7 (amino acids 50-750; UniprotKB: Q14999) were expressed and purified in-house. Coding sequences of each protein were cloned into the pET-28a(+) expression vector via NcoI and XhoI restriction sites, incorporating an N-terminal His-tag for purification. The resulting plasmids were transformed into *E. coli* BL21(DE3) competent cells. Bacterial cultures were grown at 37 °C until the optical density at 600 nm (OD) reached approximately 0.5-0.6, followed by induction with 0.2 mM IPTG. Protein expression was carried out at either 16 °C or 30 °C for 15-18 hours. Recombinant proteins were purified using Ni-NTA affinity chromatography. The elution buffer was TBS buffer (20 mM Tris-HCl, 500 mM NaCl, pH 8.0) supplemented with 250 mM imidazole. After purification, the protein was dialysed and maintained in the TBS buffer, aliquoted, and snap frozen at −80 °C for storage. Purity of each protein exceeded 85%, as assessed by SDS-PAGE.

#### In vitro assay

A portion of each reaction mixture was directly subjected to SDS-PAGE to assess cleavage efficiency. The remaining portion was used for peptide identification by mass spectrometry. After electrophoresis, the target protein bands were excised from the Coomassie-stained gels and subjected to in-gel digestion. The gel pieces were sequentially destained with 50 mM NH4HCO_3_ containing 50% (v/v) acetonitrile, dehydrated with 100% acetonitrile, reduced with 10 mM DTT at 37 °C for 60 min, and alkylated with 55 mM iodoacetamide at room temperature in the dark for 45 min. Proteins were then digested overnight with trypsin (10 ng/μL) at 37 °C. The resulting peptides were sequentially extracted with 50% acetonitrile/5% formic acid and 100% acetonitrile, followed by vacuum drying. For desalting and concentration, peptides were purified using C18 tips, eluted with 50% acetonitrile containing 0.1% trifluoroacetic acid, and vacuum-dried prior to LC-MS/MS analysis.

#### LC-MS/MS analysis

Peptides were resuspended in solvent A (0.1% formic acid and 2% acetonitrile in water) and separated using a Thermo Fisher EASY-nLC 1000 ultrahigh-performance liquid chromatography (UHPLC) system. Chromatographic separation was performed on a C18 reverse-phase analytical column (3.0 μm particle size, 75 μm inner diameter x 15 cm length). Solvent A consisted of 0.1% formic acid and 2% acetonitrile in water, while solvent B consisted of 0.1% formic acid and 98% acetonitrile. The gradient was programmed as follows: 6% to 25% solvent B over 0-16 minutes, 25% to 40% B over 16-22 minutes, 40% to 80% B over 22-26 minutes, and held at 80% B from 26-30 minutes. The flow rate was maintained at 350 nL/min. The separated peptides were ionised by a nanospray ionisation (NSI) source and introduced into a Thermo Fisher Orbitrap Fusion Lumos mass spectrometer for tandem mass spectrometry (MS/MS) analysis. The ionisation voltage was set to 2.1 kV. Both MS1 (full scan) and MS2 (fragmentation scan) spectra were acquired in the Orbitrap analyser. The MS1 scan range was 350-1800 m/z with a resolution of 70,000. The MS2 scan range started at 100 m/z, with a resolution of 17,500. Data were acquired in data-dependent acquisition (DDA) mode. The top 20 most intense precursor ions from each full scan were selected for fragmentation via higher-energy collisional dissociation (HCD) at a normalised collision energy of 28%. Dynamic exclusion was set to 15 seconds to avoid repeated fragmentation of the same precursor. The automatic gain control (AGC) target was set to 5 × 104, the signal intensity threshold was 5000 ions/s, and the maximum injection time was 200 ms.

#### LC-MS/MS data analysis

The raw LC-MS/MS data were processed and searched using PEAKS Studio 12.5. The data were searched against the Bacillus subtilis protein database. The following parameters were used for the database search: enzyme specificity was defined as Trypsin/P and Caspase3/D with up to two missed cleavages allowed; precursor mass tolerance was set to 10 ppm and fragment mass tolerance to 0.02 Da. Carbamidomethylation of cysteine was specified as a fixed modification, while oxidation of methionine and N-terminal acetylation were set as variable modifications.

## Data Availability

The benchmark dataset is collected from MEROPS (https://www.ebi.ac.uk/merops/). The predicted structures are downloaded from the AlphaFold Protein Structure Database (https://www.alphafold.ebi.ac.uk/). Source data are provided with this paper.

## Code availability

The source code of OmniCleave is publicly available at the GitHub repository (https://github.com/ABILiLab/OmniCleave). To facilitate broader use, we additionally provide a user-friendly graphical interface, comprehensive usage documentation, and a freely accessible web server at https://omnicleave.biotools.bio/.

## Acknowledgments

This work was supported by the National Natural Science Foundation of China [62202388]; the National Key Research and Development Program of China [2022YFF1000100]; and the Qin Chuangyuan Innovation and Entrepreneurship Talent Project [QCYRCXM-2022-230]. F.L. is supported by the Australian National Health and Medical Research Council (NHMRC) Investigator Fellowship (GNT2041439).

## Extended Figures

**Extended Figure 2:**
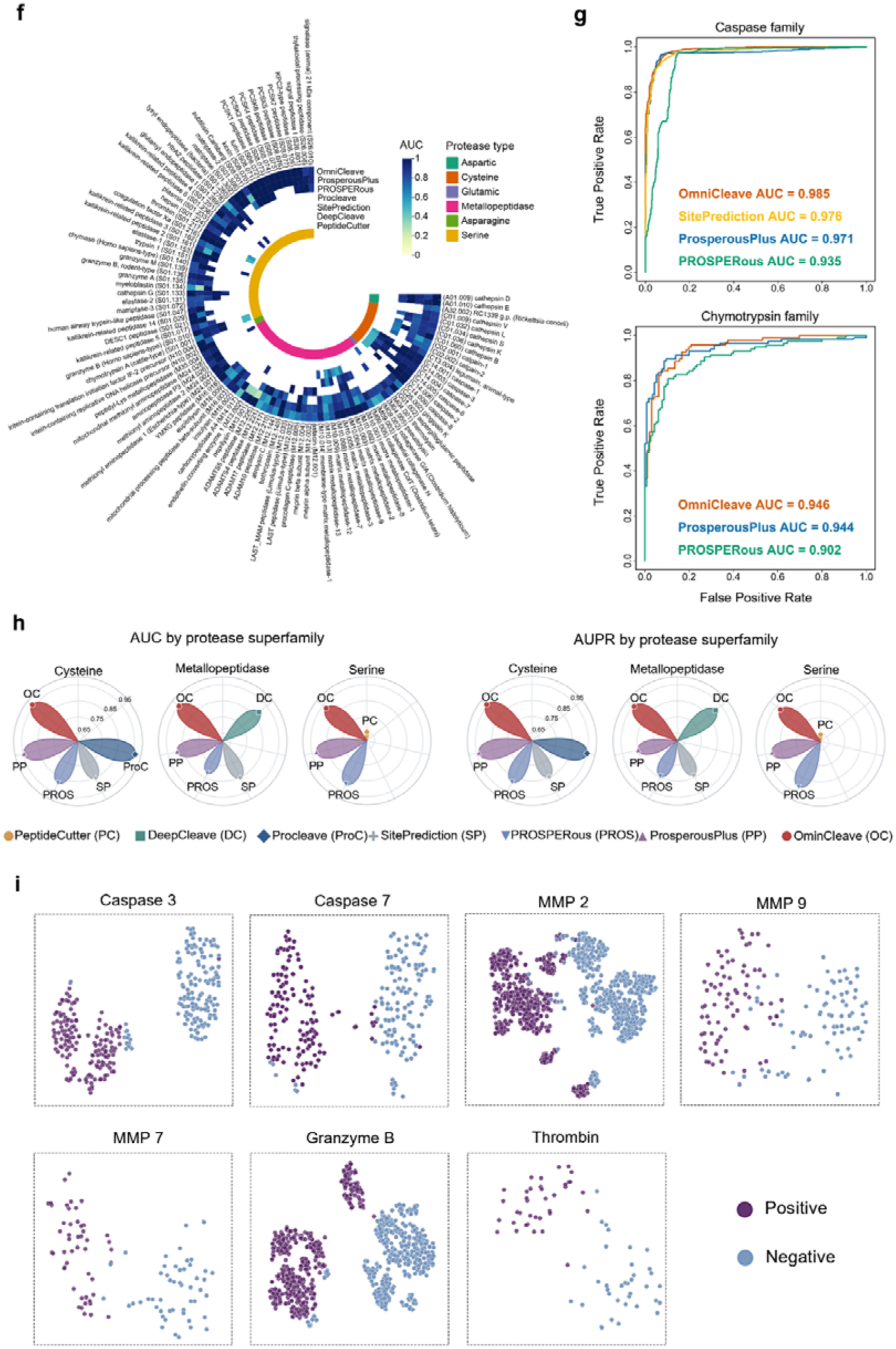
**f**. Comparison of AUC values across 103 protease substrate cleavage datasets between OmniCleave and six representative state-of-the-art tools (sequence similarity threshold: >30%). **g**. ROC curves of OmniCleave and baseline methods across the Caspase and Chymotrypsin protease families. **h**. Comparison of OmniCleave with other tools across three superfamilies (Cysteine, Metallopeptidase, and Serine). The other three superfamilies (Aspartic, Glutamic, and Asparagine) were not included in this cross-tool comparison due to the smaller number of proteases they contain and the limited number of proteases of comparative tools supported. **i**. t-SNE visualization of the fully connected layer representations of OmniCleave, shown for seven proteases (caspase-3, caspase-7, MMP2, MMP9, MMP7, granzyme B, and thrombin) in the test set.

**Extended Figure 2:**
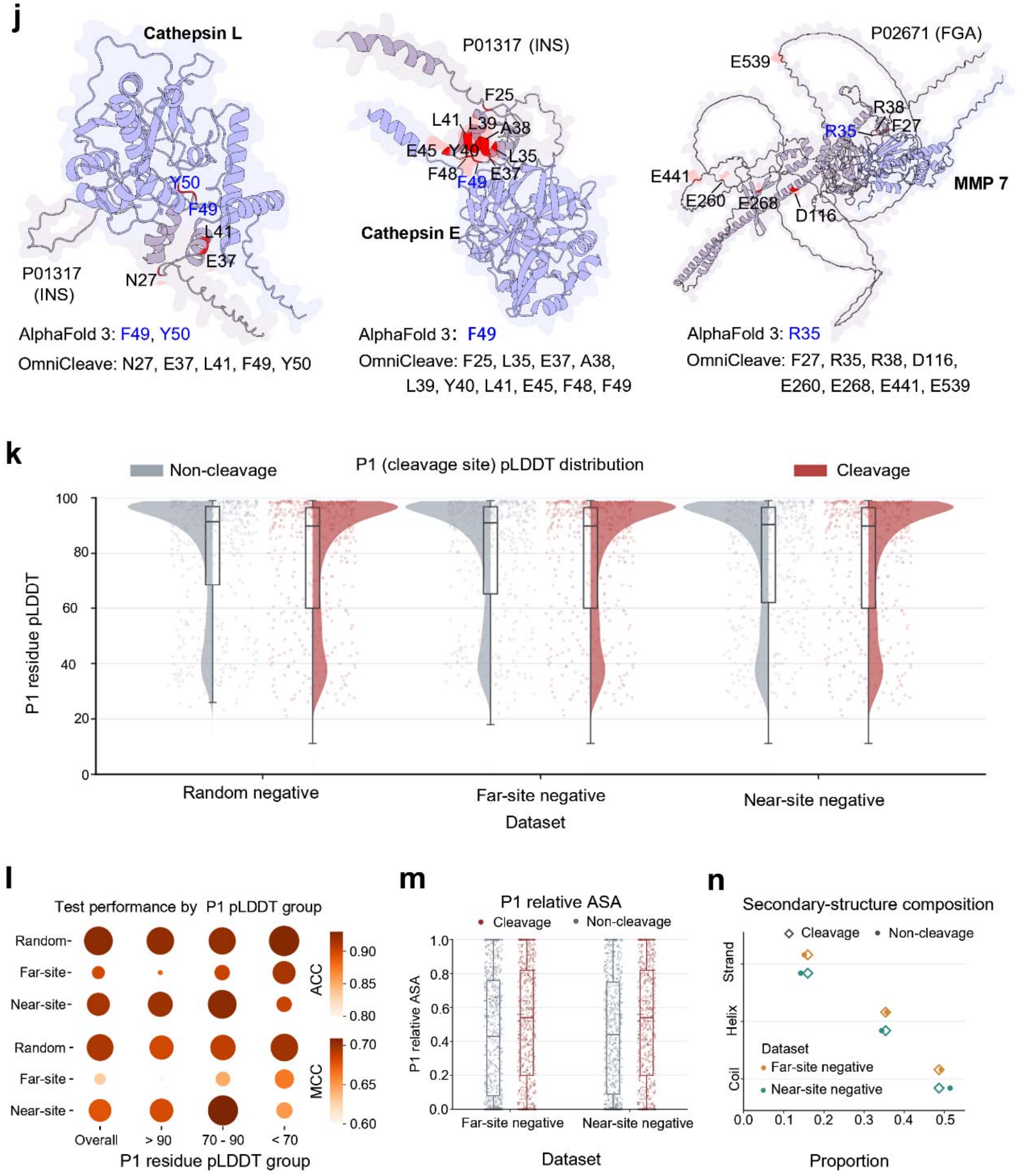
**j**. Case studies of protease-substrate complexes comparing cleavage-site predictions generated by OmniCleave and AlphaFold3. **k**. Comparison of the P1-residue pLDDT distributions between cleavage and non-cleavage sites (putative negative sites) under different negative-sampling strategies. **l**. Predictive performance of OmniCleave across different P1-residue pLDDT groups. **m**. Comparison of the ASA distributions at the P1 site for cleavage sites and non-cleavage sites under two putative negative sampling strategies. **n**. Comparison of the secondary structure distribution at the P1 site between cleavage sites and non-cleavage sites under two negative sampling strategies.

**Extended Figure 4:**
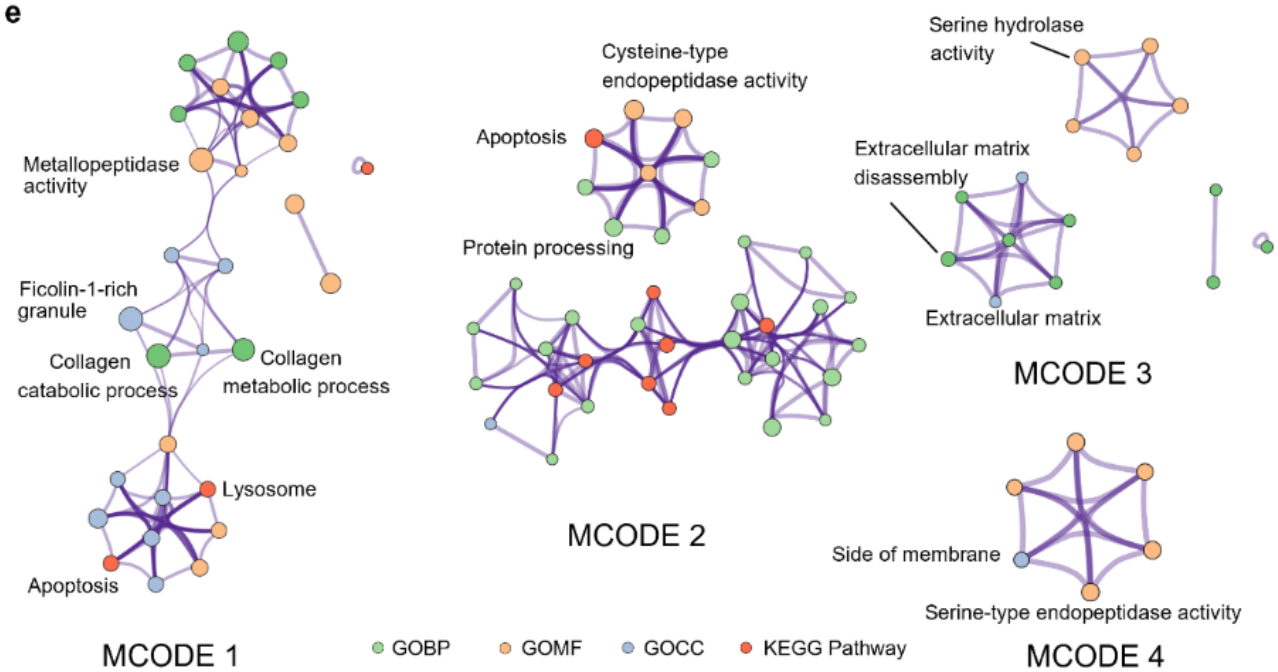
**e**. GO and KEGG enrichment analysis of proteases within MCODE clusters.

**Extended Figure 5:**
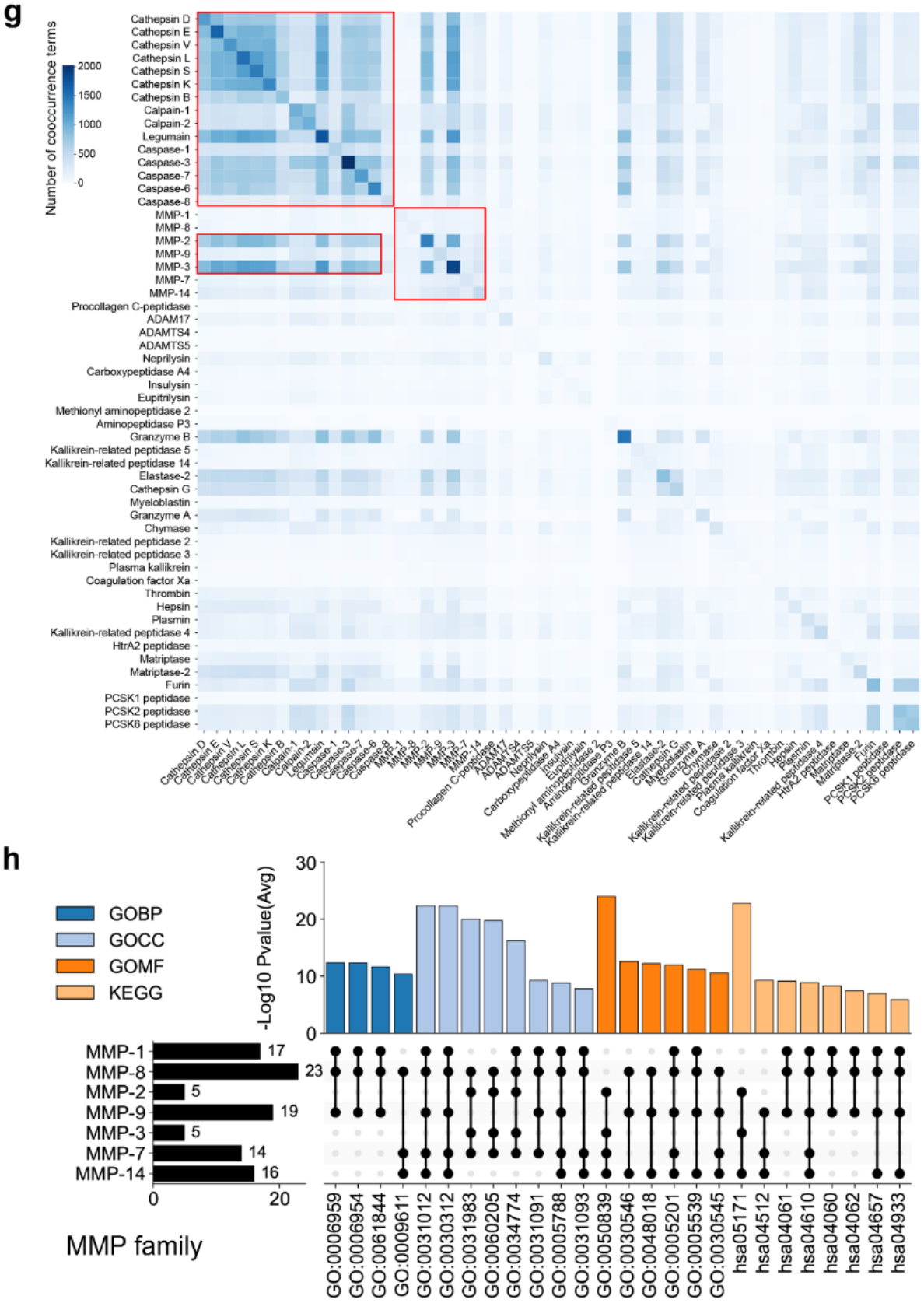
**g**. Co-occurrence enrichment terms for substrates of 54 proteases. **h**. Distribution of shared enrichment terms for substrates of the MMP family.

## Notes

### Competing Interest Statement

The authors have declared no competing interest.

### Summary of Updates

Update title Included many benchmark tests

## References

1. R. W. King, R. J. Deshaies, J.-M. Peters, M. W. Kirschner, How proteolysis drives the cell cycle. Science 274, 1652–1659 (1996).

2. V. M. Dixit, The road to death: Caspases, cleavage, and pores. Science Advances 9, eadi2011 (2023).

3. C. López-Otín, L. M. Matrisian, Emerging roles of proteases in tumour suppression. Nature reviews cancer 7, 800–808 (2007).

4. C. Hu et al., Craspase is a CRISPR RNA-guided, RNA-activated protease. Science 377, 1278–1285 (2022).

5. L. Hedstrom, Serine protease mechanism and specificity. Chemical reviews 102, 4501–4524 (2002).

6. T. R. Blum et al., Phage-assisted evolution of botulinum neurotoxin proteases with reprogrammed specificity. Science 371, 803–810 (2021).

7. B. Turk, Targeting proteases: successes, failures and future prospects. Nature reviews Drug discovery 5, 785–799 (2006).

8. C. J. Peach, L. E. Edgington-Mitchell, N. W. Bunnett, B. L. Schmidt, Protease-activated receptors in health and disease. Physiological Reviews 103, 717–785 (2023).

9. O. Schilling, C. M. Overall, Proteome-derived, database-searchable peptide libraries for identifying protease cleavage sites. Nature biotechnology 26, 685–694 (2008).

10. K. Gevaert et al., Exploring proteomes and analyzing protein processing by mass spectrometric identification of sorted N-terminal peptides. Nature biotechnology 21, 566–569 (2003).

11. Anonymous, UniProt: the universal protein knowledgebase in 2025. Nucleic acids research 53, D609–D617 (2025).

12. F. Li et al., Twenty years of bioinformatics research for protease-specific substrate and cleavage site prediction: a comprehensive revisit and benchmarking of existing methods. Briefings in bioinformatics 20, 2150–2166 (2019).

13. S. E. Boyd, R. N. Pike, G. B. Rudy, J. C. Whisstock, M. G. de La Banda, PoPS: a computational tool for modeling and predicting protease specificity. Journal of bioinformatics and computational biology 3, 551–585 (2005).

14. J. Verspurten, K. Gevaert, W. Declercq, P. Vandenabeele, SitePredicting the cleavage of proteinase substrates. Trends in biochemical sciences 34, 319–323 (2009).

15. E. Gasteiger et al., “Protein identification and analysis tools on the ExPASy server” in The proteomics protocols handbook. (Springer, 2005), pp. 571–607.

16. Z. Liu et al., GPS-CCD: a novel computational program for the prediction of calpain cleavage sites. PLoS One 6, e19001 (2011).

17. L. Hu et al., Effectively predicting HIV-1 protease cleavage sites by using an ensemble learning approach. BMC bioinformatics 23, 447 (2022).

18. A. Dumitrescu et al., TSignal: a transformer model for signal peptide prediction. Bioinformatics 39, i347–i356 (2023).

19. J. Song et al., Cascleave: towards more accurate prediction of caspase substrate cleavage sites. Bioinformatics 26, 752–760 (2010).

20. M. Wang et al., Cascleave 2.0, a new approach for predicting caspase and granzyme cleavage targets. Bioinformatics 30, 71–80 (2014).

21. J. Song et al., PROSPER: an integrated feature-based tool for predicting protease substrate cleavage sites. PloS one 7, e50300 (2012).

22. S. Kumar, B. I. Ratnikov, M. D. Kazanov, J. W. Smith, P. Cieplak, CleavPredict: a platform for reasoning about matrix metalloproteinases proteolytic events. PLoS One 10, e0127877 (2015).

23. J. Song et al., iProt-Sub: a comprehensive package for accurately mapping and predicting protease-specific substrates and cleavage sites. Briefings in bioinformatics 20, 638–658 (2019).

24. J. Song et al., PROSPERous: high-throughput prediction of substrate cleavage sites for 90 proteases with improved accuracy. Bioinformatics 34, 684–687 (2018).

25. M. Piippo, N. Lietzén, O. S. Nevalainen, J. Salmi, T. A. Nyman, Pripper: prediction of caspase cleavage sites from whole proteomes. BMC bioinformatics 11, 320 (2010).

26. F. Li et al., DeepCleave: a deep learning predictor for caspase and matrix metalloprotease substrates and cleavage sites. Bioinformatics 36, 1057–1065 (2020).

27. F. Li et al., Procleave: predicting protease-specific substrate cleavage sites by combining sequence and structural information. Genomics, proteomics & bioinformatics 18, 52–64 (2020).

28. F. Li et al., ProsperousPlus: a one-stop and comprehensive platform for accurate protease-specific substrate cleavage prediction and machine-learning model construction. Briefings in Bioinformatics 24, bbad372 (2023).

29. C. Lu et al., Prediction and design of protease enzyme specificity using a structure-aware graph convolutional network. Proceedings of the National Academy of Sciences 120, e2303590120 (2023).

30. Y. Shin, J.-K. Lee, In silico framework about prediction of collagen-derived matrikine generation and its ligand function binding to integrin for malignancy of colon cancer. In Silico Pharmacology 13, 127 (2025).

31. A. Chang et al., Homotypic fibrillization of TMEM106B across diverse neurodegenerative diseases. Cell 185, 1346–1355. e1315 (2022).

32. C. López-Otín, J. S. Bond, Proteases: multifunctional enzymes in life and disease. Journal of Biological Chemistry 283, 30433–30437 (2008).

33. T. Pan et al., Inferring transporter specificity through language model–guided graph learning. (2026).

34. B. Shi et al., DesignMaster: a multi-conditional diffusion framework for rational PROTAC design. Bioinformatics 42 (2026).

35. J. Liu, M. J. Roy, L. Isbel, F. Li, PROTAC-targeted degradation prediction with DegradeMaster. Bioinformatics 41, i342–i351 (2025).

36. K. Jha, S. Saha, H. Singh, Prediction of protein–protein interaction using graph neural networks. Scientific Reports 12, 8360 (2022).

37. J. Liu et al., MolX: A Geometric Foundation Model for Protein–Ligand Modelling. bioRxiv, 2026.2002. 2026.708362 (2026).

38. Z. Ran et al., A scalable equivariant graph network framework for precise protein function prediction. Genome Biology 26, 407 (2025).

39. N. D. Rawlings, A. J. Barrett, R. Finn, Twenty years of the MEROPS database of proteolytic enzymes, their substrates and inhibitors. Nucleic acids research 44, D343–D350 (2016).

40. U. Consortium, UniProt: a worldwide hub of protein knowledge. Nucleic acids research 47, D506–D515 (2019).

41. J. Jumper et al., Highly accurate protein structure prediction with AlphaFold. Nature 596, 583–589 (2021).

42. M. Varadi et al., AlphaFold Protein Structure Database: massively expanding the structural coverage of protein-sequence space with high-accuracy models. Nucleic acids research 50, D439–D444 (2022).

43. Z. Lin et al., Evolutionary-scale prediction of atomic-level protein structure with a language model. Science 379, 1123–1130 (2023).

44. D. Szklarczyk et al., The STRING database in 2023: protein–protein association networks and functional enrichment analyses for any sequenced genome of interest. Nucleic acids research 51, D638–D646 (2023).

45. S. Hubbard, S. Campbell, J. Thornton, Molecular recognition: conformational analysis of limited proteolytic sites and serine proteinase protein inhibitors. Journal of molecular biology 220, 507–530 (1991).

46. Y. Zhou et al., Metascape provides a biologist-oriented resource for the analysis of systems-level datasets. Nature communications 10, 1523 (2019).

47. P. Shannon et al., Cytoscape: a software environment for integrated models of biomolecular interaction networks. Genome research 13, 2498–2504 (2003).

48. G. D. Bader, C. W. Hogue, An automated method for finding molecular complexes in large protein interaction networks. BMC bioinformatics 4, 2 (2003).

49. R. F. Alford et al., The Rosetta all-atom energy function for macromolecular modeling and design. Journal of chemical theory and computation 13, 3031 (2017).

50. D. Stickens et al., Altered endochondral bone development in matrix metalloproteinase 13-deficient mice. (2004).

51. J. Oh et al., Mutations in two matrix metalloproteinase genes, MMP-2 and MT1-MMP, are synthetic lethal in mice. Oncogene 23, 5041–5048 (2004).

52. A. Ducharme et al., Targeted deletion of matrix metalloproteinase-9 attenuates left ventricular enlargement and collagen accumulation after experimental myocardial infarction. The Journal of clinical investigation 106, 55–62 (2000).

53. L. A. Rudolph-Owen, D. L. Hulboy, C. L. Wilson, J. Mudgett, L. M. Matrisian, Coordinate expression of matrix metalloproteinase family members in the uterus of normal, matrilysin-deficient, and stromelysin-1-deficient mice. Endocrinology 138, 4902–4911 (1997).

54. Z. Li, M. Sheng, Caspases in synaptic plasticity. Molecular brain 5, 15 (2012).

55. J.-Y. Wang, Z.-G. Luo, Non-apoptotic role of caspase-3 in synapse refinement. Neuroscience bulletin 30, 667–670 (2014).

56. M. d’Amelio et al., Caspase-3 triggers early synaptic dysfunction in a mouse model of Alzheimer’s disease. Nature neuroscience 14, 69–76 (2011).

57. D. Chéreau, L. Kodandapani, K. J. Tomaselli, A. P. Spada, J. C. Wu, Structural and functional analysis of caspase active sites. Biochemistry 42, 4151–4160 (2003).

58. M. Sulpizi, U. Rothlisberger, P. Carloni, Molecular dynamics studies of caspase-3. Biophysical journal 84, 2207–2215 (2003).

59. P. Schake, S. N. Bolz, K. Linnemann, M. Schroeder, PLIP 2025: introducing protein–protein interactions to the protein–ligand interaction profiler. Nucleic Acids Research 53, W463–W465 (2025).

60. W. L. DeLano, Pymol: An open-source molecular graphics tool. CCP4 Newsl. protein crystallogr 40, 82–92 (2002).

61. L. Fu, B. Niu, Z. Zhu, S. Wu, W. Li, CD-HIT: accelerated for clustering the next-generation sequencing data. Bioinformatics 28, 3150–3152 (2012).

62. W. Kabsch, C. Sander, Dictionary of protein secondary structure: pattern recognition of hydrogen-bonded and geometrical features. Biopolymers: Original Research on Biomolecules 22, 2577–2637 (1983).

63. X. Kong, W. Huang, Y. Liu, Generalist equivariant transformer towards 3d molecular interaction learning. arXiv preprint arXiv:2306.01474 (2023).

64. X. Glorot, Y. Bengio (2010) Understanding the difficulty of training deep feedforward neural networks. in Proceedings of the thirteenth international conference on artificial intelligence and statistics (JMLR Workshop and Conference Proceedings), pp 249–256.

65. L. Liu et al., Technical report of HelixFold3 for biomolecular structure prediction. arXiv preprint arXiv:2408.16975 (2024).

66. A. Singh, M. M. Copeland, P. J. Kundrotas, I. A. Vakser, “GRAMM web server for protein docking” in Computational drug discovery and design. (Springer, 2023), pp. 101–112.

